# MYC activity is required for maintenance of the Neuromesodermal Progenitor signalling network and for correct timing of segmentation clock gene oscillations

**DOI:** 10.1101/212035

**Authors:** Ioanna Mastromina, Laure Verrier, Kate G. Storey, J. Kim Dale

## Abstract

The Myc transcriptional regulators are implicated in a range of cellular functions, including proliferation, cell cycle progression, metabolism and pluripotency maintenance. Here, we investigated the expression, regulation and function of Myc during mouse embryonic axis elongation and segmentation. Expression of both *cMyc* and *MycN* in the domains where neuromesodermal progenitors (NMPs) and underlying caudal pre-somitic mesoderm (cPSM) cells reside is coincident WNT and FGF signals; factors known to maintain progenitors in an undifferentiated state. Pharmacological inhibition of MYC activity, downregulates expression of WNT/FGF components. In turn, we find that *cMyc* expression is WNT, FGF and NOTCH regulated, placing it centrally in the signalling circuit that operates in the tail end that both sustains progenitors and drives maturation of the PSM into somites. Interfering with MYC function in the PSM, where it displays oscillatory expression, delays the timing of segmentation clock oscillations and thus of somite formation. In summary, we identify Myc as a component that links NMP maintenance and PSM maturation during the body axis elongation stages of mouse embryogenesis.

**Summary Statement:** MYC operates in a positive feedback loop with WNT/FGF signals to maintain the progenitors which facilitate body axis elongation while its activity is crucial for timing of the segmentation clock.

## Introduction

The Myc family of proto-oncogenes has been characterised as one of the most exhaustively studied family of vertebrate genes (Meyer and Penn, 2008, Eilers and Eisenman, 2008). Since the discovery of cMyc (Alitalo et al., 1983, Watson et al., 1983), two more family members were identified, namely MycN (Brodeur et al., 1984, Emanuel et al., 1985)and L-Myc (Nau et al., 1985, Ikegaki et al., 1989), and a plethora of studies has now placed each member centrally in cancer initiation and progression, in a context specific manner (reviewed in Tansey (2014)). It is now established that the oncogenic potential of Myc is mediated through the transcriptional control of multiple target gene sets (Dang et al., 2006, Zeller et al., 2003, Zeller et al., 2006). Myc contains a basic helix-loop-helix (bHLH) domain and transcriptional activation takes place when it heterodimerises with its obligate partner Max (Blackwood and Eisenman, 1991, Blackwood et al., 1991), while it can act as a repressor when it dimerises with Miz1 (Staller et al., 2001). Additional co-factors, such as the bromodomain containing protein BRD4, mediate recruitment of the Myc complex onto the chromatin.

The discovery of cMyc as one of the four Yamanaka factors required for the reprogramming of adult terminally differentiated cells back into the pluripotent cell state (Takahashi and Yamanaka, 2006) has refocused Myc-related research on stem cell biology and has highlighted multiple roles for Myc including regulation of proliferation, cell cycle and metabolism, within the pluripotent cell state (reviewed in Fagnocchi and Zippo (2017). In parallel, recent studies have highlighted diverse roles for Myc during embryo development, including regulation of metabolism of the pre-implantation embryo (Scognamiglio et al., 2016) progenitor sorting and fitness through the process of cell competition upon differentiation onset in the early post-implantation epiblast (Claveria et al., 2013, Sancho et al., 2013), maintenance of the size of the neural crest progenitor pool (Kerosuo and Bronner, 2016) and neural differentiation progression in the chicken embryo neural tube (Zinin et al., 2014).

Both cMyc and MycN homozygote mutant mice are embryonic lethal by mid gestation displaying a range of defects (Davis et al., 1993, Trumpp et al., 2001, Sawai et al., 1993) suggesting that the multifunctional Myc factors hold important roles during development, and likely in a context specific manner. Expression pattern analyses have highlighted the presence of both *cMyc* and *MycN* in multiple embryonic tissues (Downs et al., 1989, Kato et al., 1991, Ma et al., 2014). However, these data, based on radiolabelled probes, give very low definition and low signal to noise ratio, and as such cannot be utilised to decipher precise patterns of expression for the Myc members. For example, detailed expression pattern and specific functions of the Myc genes during elongation and segmentation of the embryo body axis, has yet to be investigated with respect to the different progenitor subpopulations that comprise the tail region at these developmental stages (Wymeersch et al., 2016). In particular, it is now established that the E8.5 post-implantation epiblast is a heterogeneous domain where progenitors with different developmental potentials reside (Wilson et al., 2009, Wymeersch et al., 2016, Henrique et al., 2015).

Key to this study, detailed fate mapping and clonal analysis has indicated that posterior neural and mesoderm lineages emerge from a common progenitor population, termed Neuromesodermal progenitors (NMPs) (Cambray and Wilson, 2002, Cambray and Wilson, 2007, Tzouanacou et al., 2009, Delfino-Machín et al., 2005, Rodrigo Albors et al., 2016). NMPs have been identified so far in the human, mouse chicken and zebrafish embryos (Olivera-Martinez et al., 2012, Wymeersch et al., 2016, Goto et al., 2017) and have been generated *in vitro* from both mouse and human ESC (Gouti et al., 2017, Gouti et al., 2014, Tsakiridis et al., 2014, Turner et al., 2014, Verrier et al., 2017). In the mouse embryo, NMPs first arise at E7.5, in the domain of Node Streak Border (NSB) and associated caudal-lateral epiblast (CLE), persist in the NSB and CLE at E8.5 and are subsequently incorporated in the Chordo-Neural Hinge (CNH) during tail growth stages (reviewed in Henrique et al. (2015)). Importantly, the dual-fated NMPs supply cells both to the forming neural plate (open pre-neural tube) and to the caudal presomitic mesoderm (cPSM) (Tzouanacou et al., 2009, Rodrigo Albors et al., 2016, Gouti et al., 2014), which further matures and segments rostrally to form the somites on either side of the neural tube. Throughout this process, the NMPs and cPSM cells are maintained in an “undifferentiated” progenitor state by the activity of two main signalling pathways, WNT and FGF, components of which show very high expression in the posterior of the embryo (Wilson et al., 2009, Hubaud and Pourquie, 2014). In addition, WNT, FGF and NOTCH signalling pathways comprise the molecular machinery of the segmentation clock, a molecular oscillator which regulates the periodic segmentation of the PSM into somites (Maroto et al., 2012, Hubaud and Pourquie, 2014). Concomitantly, neural and somitic differentiation is promoted by high levels of Retinoic Acid (RA), which is produced by the forming somites and counteracts WNT/FGF signalling (Delfino-Machín et al., 2005, Diez del Corral et al., 2003, Olivera-Martinez and Storey, 2007, Dequeant and Pourquie, 2008, Dubrulle and Pourquie, 2004, Naiche et al., 2011, Sakai et al., 2001). Interestingly, *cMyc* has been shown to be present and display dynamic oscillatory mRNA expression in the mouse and chick pre-somitic mesoderm (Dequeant et al., 2006, Krol et al., 2011), while also being expressed in high levels in the domain that harbours the NMPs in the chicken embryo (Olivera-Martinez et al., 2014). However, no investigation as to the functional significance of Myc expression in these domains has been conducted.

Here we elucidate divergent roles for Myc during posterior embryonic body axis formation. We find that cMyc is indispensable for the proper timing of clock gene oscillations through regulation of NOTCH signalling. Moreover we demonstrate Myc operates in a positive feedback loop with WNT and FGF signalling in the CLE of the E8.5 embryo and that inhibition of MYC activity results in transcriptional downregulation of different gene sets, which include regulators of metabolism. These findings are the first to provide a common regulator of different sets of genes that co-ordinate progenitor cell maintenance, metabolism and differentiation in the NMPs and cPSM in the mouse embryo.

## Results

### *cMyc* is expressed in the CLE and underlying pre-somitic mesoderm and its expression persists during axial elongation and body axis segmentation

We generated *cMyc* and *MycN* riboprobes and carried out an initial expression pattern analysis (Figure 1). Both genes show high expression in the head structures and in particular *cMyc* is expressed in high levels in the somites and in lower levels in the neural tube (Figure 1A). In contrast, *MycN* shows high levels in the neural tube and low levels in somitic tissue (Figure 1B). We were particularly interested to see that both Myc members show high levels of expression in the posterior of the embryo proper. In particular we find high levels of *cMyc* in the CLE domain, and lower levels in the underlying cPSM (Figure 1B a-a’). *MycN* exhibits a converse expression pattern, low in CLE and higher in the underlying cPSM (Figure 1B b-b’). The CLE is the region where a small bipotent population of precursors is located, namely the neuromesodermal progenitors (NMPs). These cells can be visualised by the co-expression of Sox2 and Brachyury (Figure 1A) and maintenance of their bipotency relies on autocrine and paracrine WNT/FGF signalling. We find that both *Myc* factors are expressed alongside *Wnt3a* and *Fgf8* in the CLE and alongside the NOTCH target gene *Lfringe* (Dale et al., 2003, McGrew et al., 1998) in *the* cPSM (Figure 1B). Using immunofluorescence labelling we find that cMyc is co-expressed with Sox2 and Brachyury in the CLE and underlying cPSM (Figure 1C). These expression data therefore show that mouse NMPs co-express cMyc, *Wnt3a*, *Fgf8*, Sox2 and Brachyury. Crosstalk between cMyc and FGF (Yu et al., 2017) or WNT (Fagnocchi et al., 2016) or Sox2 (Lin et al., 2009) has been reported in other systems. It is therefore likely that cMyc might be involved in the NMP signalling network.

**Figure 1:**
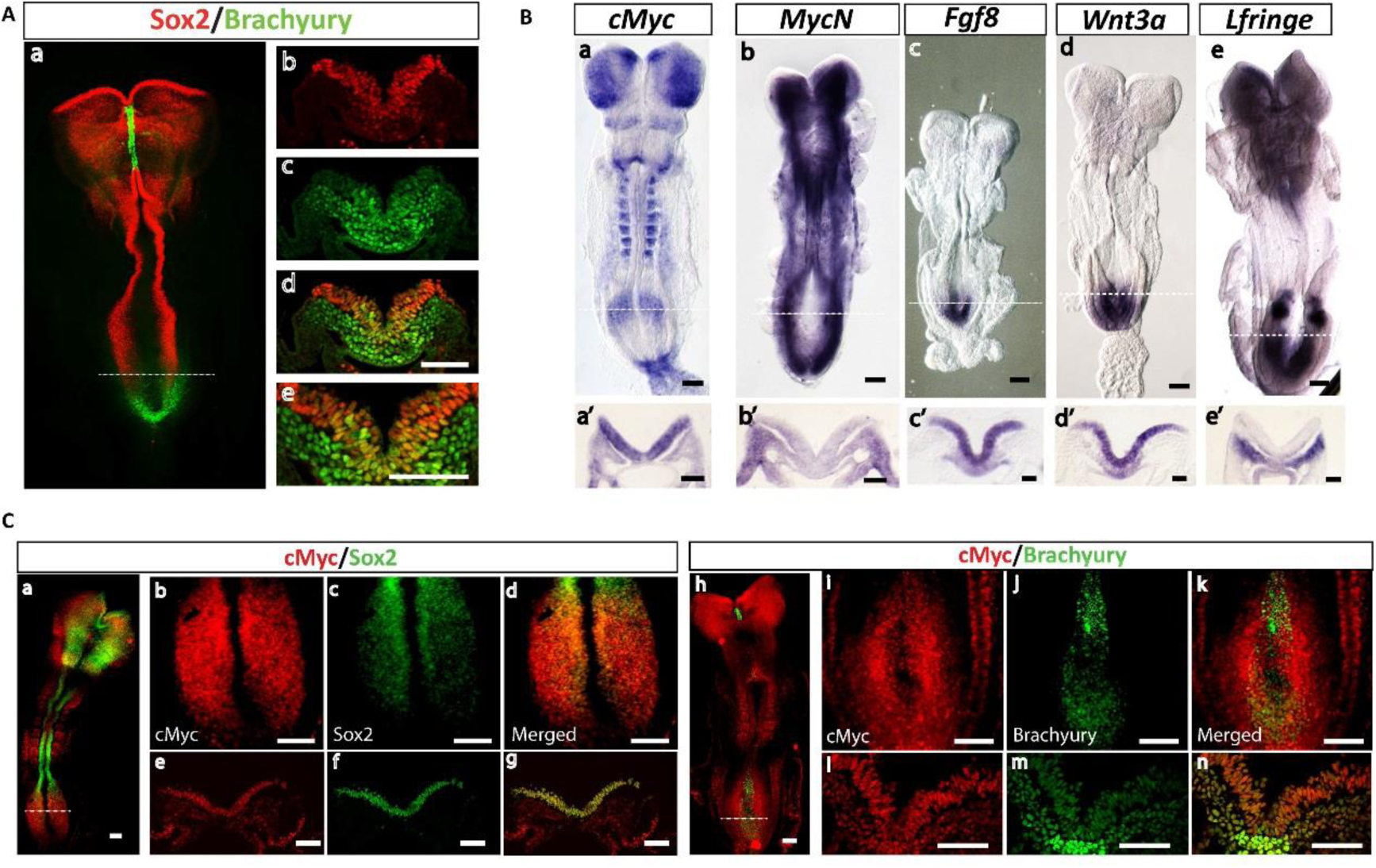
cMyc is co-expressed with *Wnt3a*, *Fgf8*, Sox2 and Brachyury in the CLE. A. Representative confocal images of an E8.5 embryo labelled by immunofluorescence for Sox2 and Brachyury (n=3 embryos). (a) Wholemount E8.5 and (b-d) are transverse sections at the level of the CLE domain (demarcated by the white dotted line in (a)). Sox2 labels the neuroepithelium along the A/P axis and the CLE whereas Brachyrury labels the PSM and tailbud mesoderm. (e) is a magnification of (d) and shows the location of the NMPs (Sox2/Brachyury co-expressing cells) in the CLE epithelium.
B. Representative *in situ* hybridisation images of E8.5 embryos stained for mRNA expression of (a) *cMyc* (n= 10 embryos) and (b) *MycN* (n=4 embryos) (c) *Fgf8* (n=3 embryos), (d) *Wnt3a* (n=3 embryos) and (e) *LFringe* (n=5 embryos). (a’-e’) are transverse sections of the CLE and underlying cPSM domain indicated by white dotted lines on (a-e). (a’) *cMyc*, (b’) *Fgf8* and (c’) *Wnt3a* show high levels of expression in the CLE. (b’) *MycN* and (e’) *LFringe* show high levels of expression in the cPSM.
C. Representative confocal images of immunofluorescence labelling of E8.5 embryos for cMyc and Sox2 (a-g; n=3 embryos) and cMyc and Brachyury (h-n; n= 3 embryos). Sox2/cMyc co-expressing cells are evident in the transverse sections of the CLE (e’-g’) panels correspond to sections at the level of the domain demarcated by the white dotted line in (a). Brachyury/cMyc co-expressing cells are evident both in CLE and underlying cPSM (l’-m’ panels, correspond to white dotted line in (e)). Scale bars are 100 μm

### cMyc is expressed in the tail bud at E9.5 and E10.5 and displays oscillatory mRNA expression in the PSM

We further characterised expression of *cMyc* and *MycN* during E9.5 and E10.5, the embryonic stages where the anterio-posterior axis elongates and segments into somites (Gibb et al., 2010, Henrique et al., 2015). The tail bud mesoderm is the main reservoir of cPSM progenitors, whereas the caudal most, Sox2/Brachyury positive region of the neuroepithelium harbours the NMPs. *Using in situ* hybridisation we find that *cMyc* displays dynamic mRNA expression in the PSM, reminiscent of clock gene expression (Figure 2A,d) consistent with previous published data in the mouse and chick PSM (Dequeant et al., 2006, Krol et al., 2011). We find that cMyc protein is expressed in the caudal most neuroepithelium (labelled by Sox2 and Brachyury) and adjacent tailbud mesoderm (labelled by Brachyury) (Figure 2B). *MycN* is expressed in the tailbud at E9.5 however its expression is downregulated at E10.5 and E11.5 (Suppl. Figure 1).

**Figure 2:**
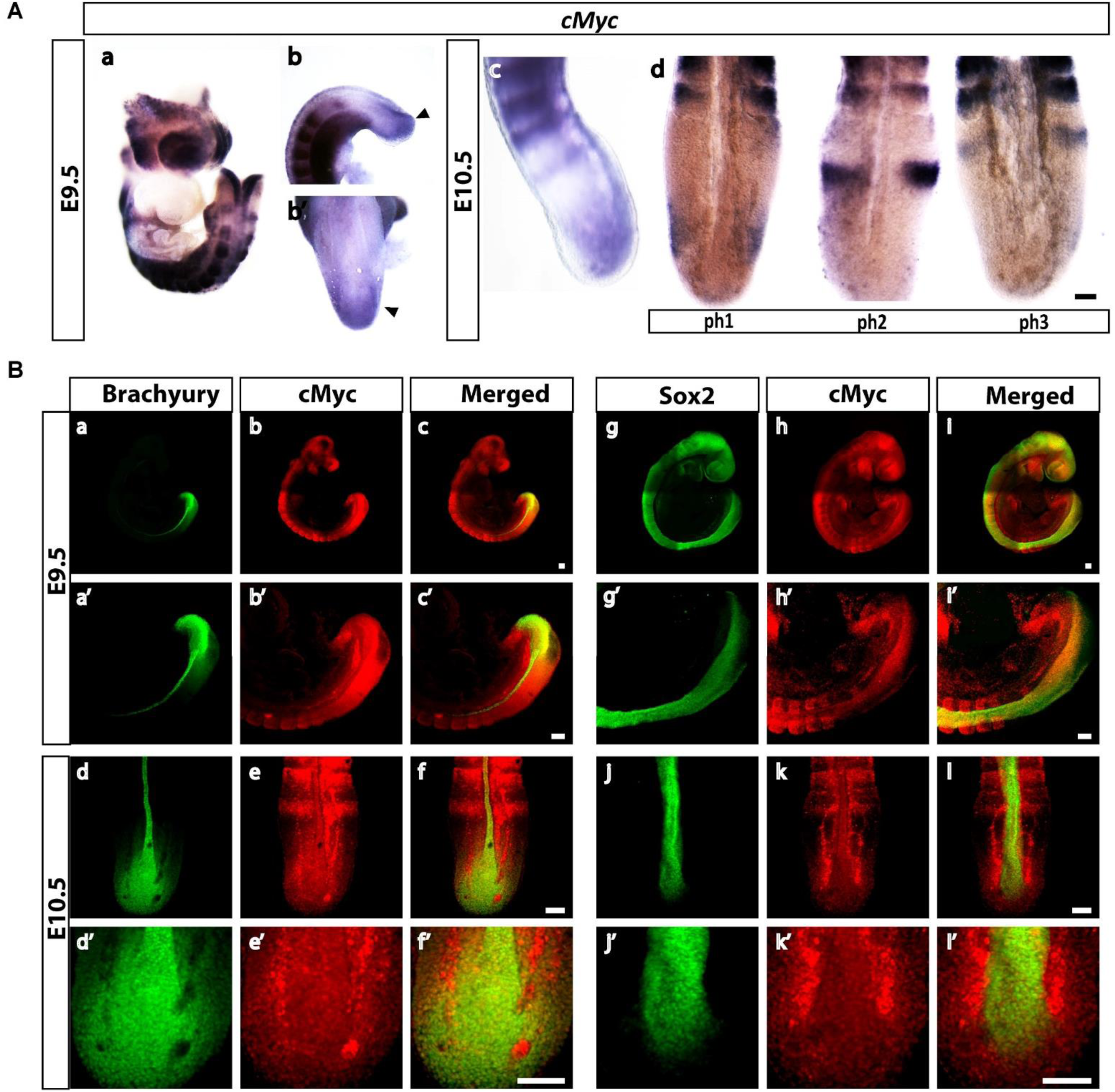
cMyc expression persists in the tail bud during E9.5-E10.5 and shows dynamic expression at the transcript level. A. Representative *in situ* hybridisation of (a) wholemount E9.5 embryo labelled for *cMyc* at E9.5 (n=12 embryos).(b-b’) high levels of cMyc are present in the caudal most neuroepithelium and adjacent PSM (black arrowheads) (c) side view of an E10.5 tail labelled for *cMyc* mRNA. (d) Three different expression profiles for *cMyc* in the PSM of E10.5 embryos, reminiscent of the three phases of the segmentation clock gene expression (n=10 embryos).
B. Representative confocal images of immunofluorescence labelling for cMyc in E9.5 and E10.5 embryos. (a-f) show cMyc and Brachyury staining in wholemount embryos. (a’-f’) are higher magnification images of (a-f) showing cMyc/Brachyury co-expressing cells in the tailbud (n=5 embryos). (g-l) show cMyc and Sox2 labelling in wholemount embryos and (g’-l’) are higher magnification images of (g-l) showing cMyc/Sox2 co-expressing cells in the tailbud. (n=5 embryos). Scale bars are 100 μm.

### Suppression of MYC activity attenuates expression of key FGF/WNT components leading to loss of the NMP dual identity

A small molecule approach was next used to investigate whether MYC activity regulates expression of key components of the WNT/FGF/NOTCH network that operates in the CLE/cPSM. To this end, we micro-dissected explant pairs that contained the NMPs and underlying cPSM from E8.5 embryos and cultured them for 6 hours in the presence of small molecule inhibitors that have been extensively used to interfere with cMyc and MycN function *in vitro* (Delmore et al., 2011, Horne et al., 2014, Posternak and Cole, 2016, Yin et al., 2003). Two different small molecules, which act via distinct molecular mechanisms, were used to cross-validate the specificity of our findings. First we used JQ1, a small molecule that competitively binds to BRD4, a co-factor that recruits the Myc complex onto the chromatin (Delmore et al., 2011) and repeated the experiments using 10074G5, a compound that interferes with heterodimerisation of Myc with its binding partner, Max (Yin et al., 2003). As a readout of small molecule efficacy we quantified, by RT-qPCR, the expression levels of two well-established Myc targets, *cyclin E1,* and *p21* (Zeller et al., 2003) and found that upon treatment with either inhibitor, *p21* levels were significantly increased whereas *cyclinE1* levels significantly decreased (Figure 3g, h). This is consistent with negative regulation of *p21* by Myc and positive Myc input on *cyclin E1* expression reported previously (Claassen and Hann, 2000, Gartel et al., 2001, Perez-Roger et al., 1997).

**Figure 3:**
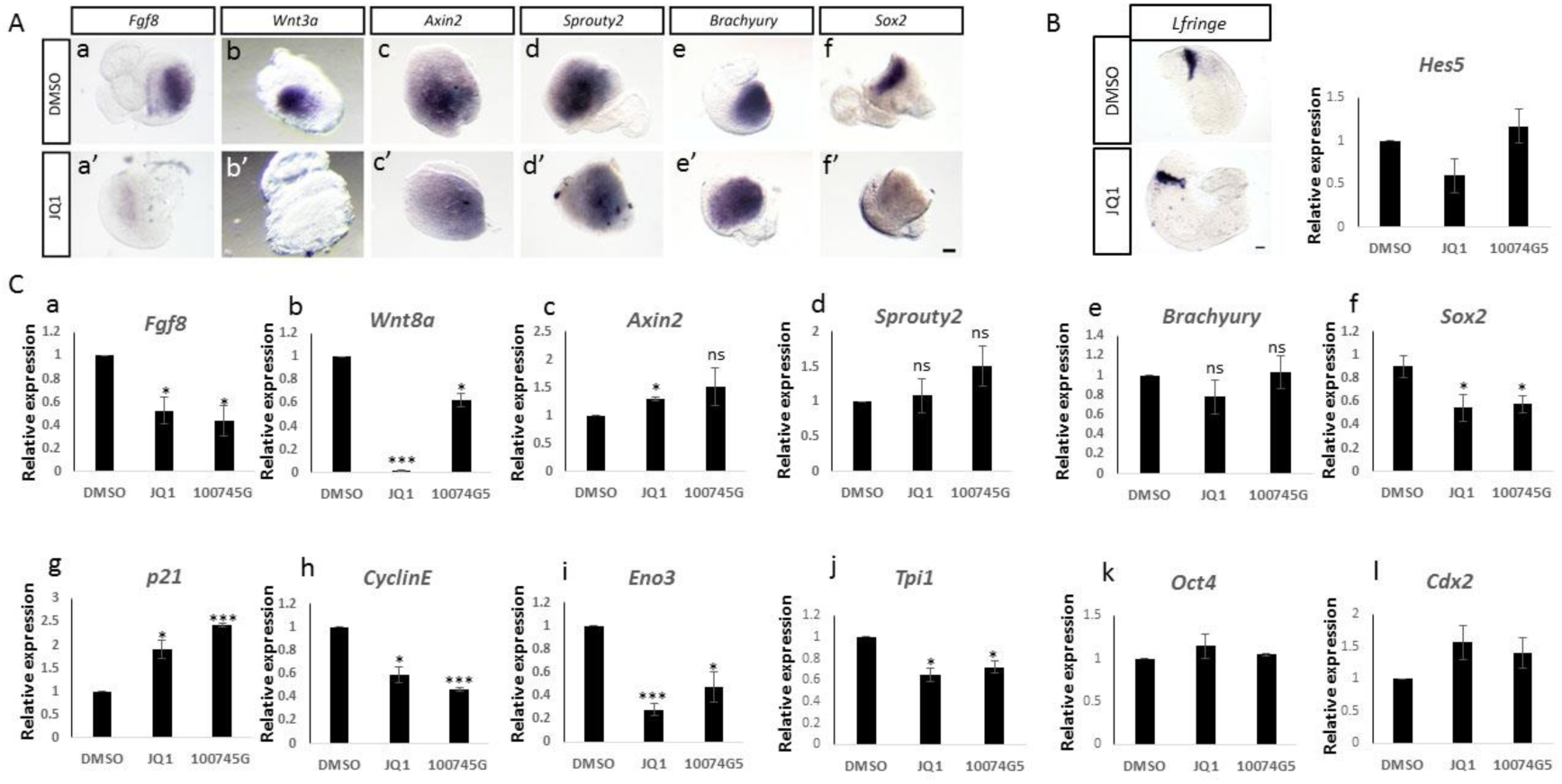
MYC activity suppression results in downregulation of *Wnt3a/8a*, *Fgf8* and *Sox2*. A. Representative *in situ* hybridisation images on CLE/cPSM explants treated with DMSO (a-f) or 10 μΜ JQ1 (a’-f’) for 6h. *Fgf8* (a-a’; n=8/8 embryos), *Wnt3a* (b-b’; 5/5 embryos) and *Sox2* (f-f’; 3/3 embryos) expression is suppressed upon JQ1 treatment. In contrast expression of *Axin2* (c-c’; n=4/4 embryos), *Sprouty2* (d-d’; n=4/4 embryos) and *Brachyury* (e-e’; n= 6/6 embryos) is not affected. Scale bar is 100 μm.
B. Representative *in situ* hybridisation images of half tail explants from E8.5 embryos (micro-dissected below the level of the last somite pair) treated either with DMSO or 10 μΜ JQ1 for 6h show no effect on expression of the NOTCH target gene *LFringe* (n=4/4 4mbryos). RT-qPCR analysis of CLE/cPSM explants for *Hes5* expression show no differences upon treatment with 10 μΜ JQ1 or 75 μΜ 10074G5 for 6h. Scale bar is 100 μm
C. Characterisation of gene expression changes in CLE/cPSM explants upon 10 μΜ JQ1 or 75 μΜ 10074G5 for 6h. Relative gene expression, normalised to actin levels. Data from 3 independent experiments, presented as Mean ± SEM.

We then assessed expression levels of key components of the WNT, FGF and NOTCH pathways known to be expressed in the tail region using RT-qPCR and *in situ* hybridisation. In explants where MYC activity was suppressed, a sharp downregulation of *Fgf8*, *Wnt3a/8a* and *Sox2* transcripts was observed, indicating that MYC activity is crucial for the integrity of the signalling circuit that operates in the NMP and cPSM progenitor cells (Figure 3Aa-b’, f, f’ Ca,b,f). Importantly, *Axin2*, *Sprouty2*, *Lfringe* and *Hes5* expression levels are unaltered at this 6h time-point revealing that despite the reduction in FGF and WNT ligand transcripts, WNT, FGF, and NOTCH target gene expression is not compromised (Figure 3 Ac-d’, B, Cc, d). In addition, even though *Sox2* expression (indicative of NMP identity in this domain) is affected in the explants, the core epiblast identity (as judged by *Cdx2*, *Oct4* mRNA expression; reviewed in Deschamps and Duboule (2017)) is not affected (Figure 3Ck,l). In addition we checked the expression levels of several metabolic genes identified recently to show high expression in the tailbud (Oginuma et al., 2017) and found that two of them, *Triosephosphate isomerase 1* (*Tpi1*) and *Enolase 3* (*Eno3*), show significant downregulation upon Myc activity suppression, consistent with Myc controlling expression of glycolytic genes in other contexts (Kim et al., 2004, Hsieh et al., 2015, Stine et al., 2015) (Figure 3Ci, j).

To further corroborate our hypothesis that Myc is important for the maintenance of WNT/FGF signalling we repeated this investigation in a pure NMP population generated *in vitro* from human Embryonic stem cells (hESC) (Verrier et al., 2017). Successful differentiation to the NMP state was verified by immunofluorescence showing co-expression of Sox2 and Brachyury and co-expressing cells could be maintained in vitro for 24h (Suppl. Figure 5). Treatment with either 500 nM JQ1 or 25 μΜ 10074G5 resulted in downregulation of *Sox2, Brachyury, Wnt8a and Fgf8*, despite the excess of WNT/FGF proteins that are present in the culture medium of the hNMPs (Figure 4). In addition we tested whether exposure to JQ1 reduces *cMyc* levels as previously reported in multiple myeloma cell lines (Delmore et al., 2011) however we did not observe changes in *cMyc* transcript levels (Figure 4). This result is in accordance with published literature which shows cMyc levels are unaffected upon JQ1 treatment in embryonic stem cells (Horne et al., 2014).

**Figure 4:**
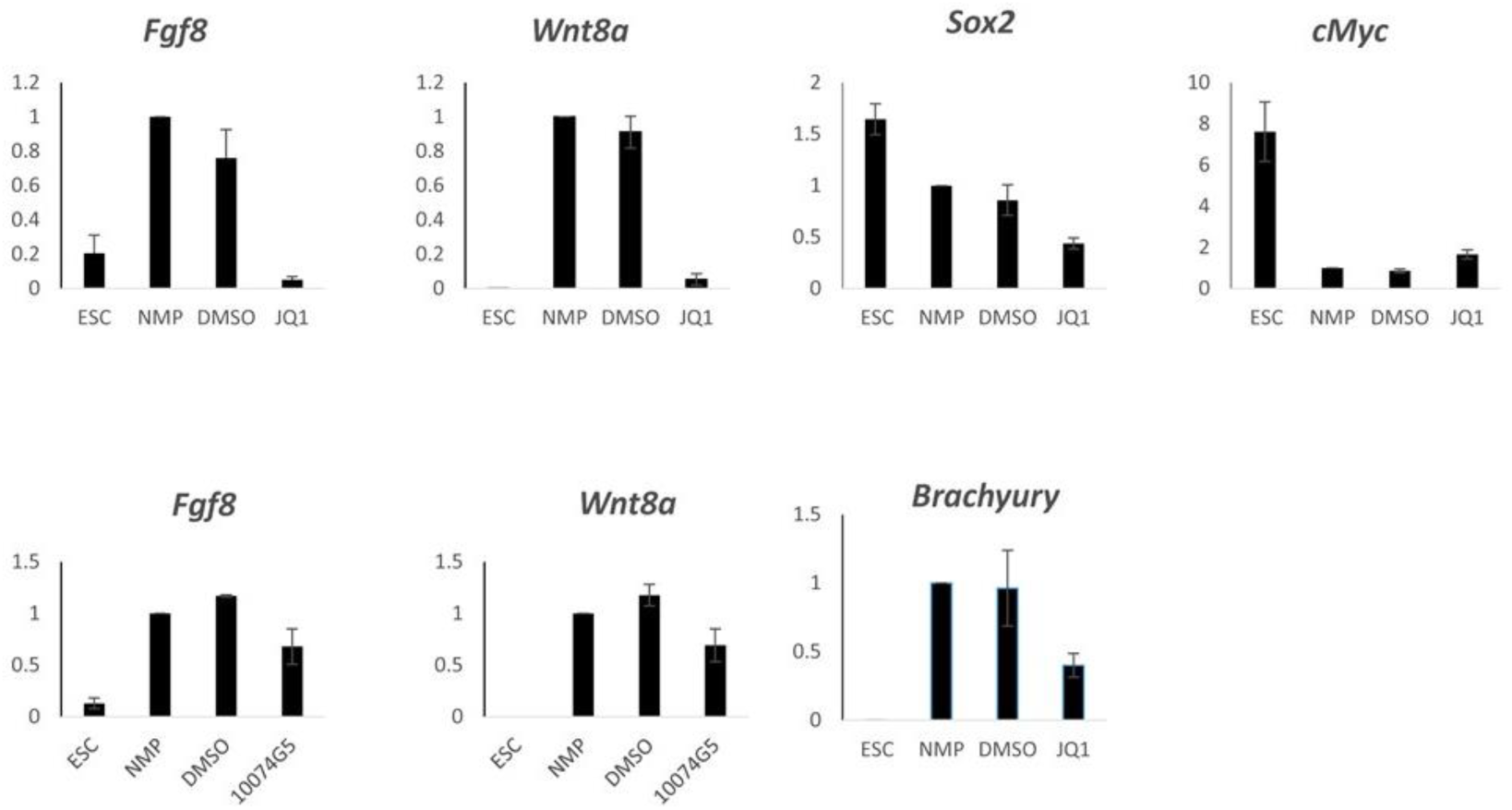
MYC activity suppression in *in vitro* hNMPS results in downregulation of *FGF8, Wnt8a, Sox2* and *Brachyury*. 500 nM of JQ1 were applied for 24h or 25 μΜ of 10074G5 were applied for 12h. Relative gene expression, normalised to expression of PRT2. Data presented as Mean ± SEM. Data from a minimum of two independent experiments

Taken together these data indicate a specific requirement for MYC dependent transcription of key NMP maintenance factors, namely *Wnt3a/8a, FGF8* and *Sox2*.

### Alleviation of Myc inhibition required for neural and mesodermal differentiation

We next investigated the possibility that transcriptional downregulation of WNT and FGF ligands, following MYC activity loss, leads to differentiation progression. Therefore culture with 10 μΜ JQ1 was increased to 10h. Neither *Pax6* (neural progenitor marker gene; (Stoykova et al., 1996)) nor *Paraxis* (rostral paraxial mesoderm marker; (Burgess et al., 1995)) expression were detected (Figure 5C). This suggests either that longer culture is required for differentiation or that MYC activity is important for initiation and/or progression of differentiation. To define better the effects of MYC inhibition after 10h we next assessed the impact of this treatment on read-outs for FGF and WNT signalling. Indeed we observed that WNT transduction is attenuated as indicated by *Axin2* transcription, however expression of the FGF target gene *Sprouty2* is not significantly affected (Figure 5E).

**Figure 5:**
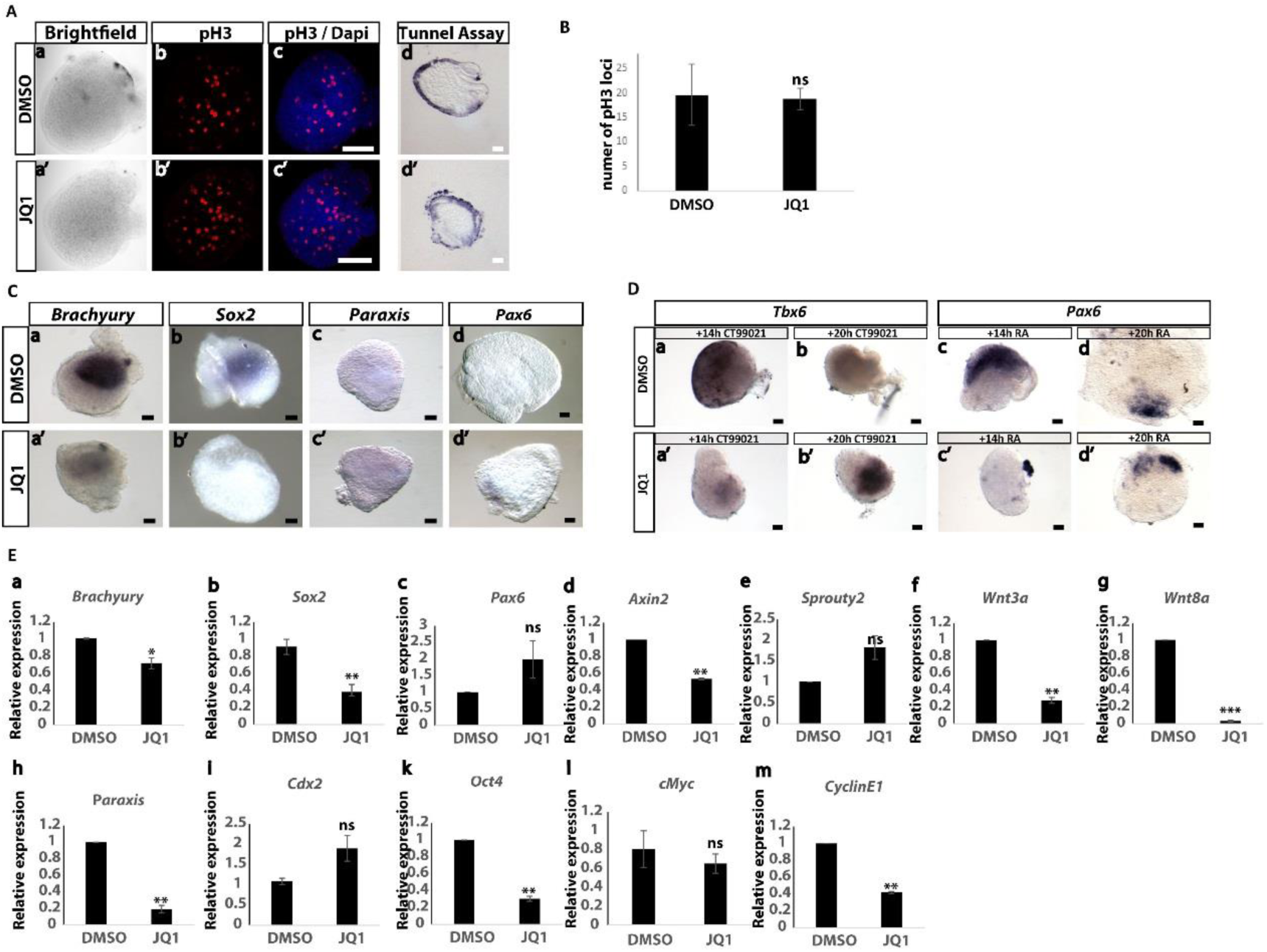
MYC activity suppression does not promote differentiation nor does it compromise viability and proliferation. A. Representative confocal images of CLE/cPSM explants cultured for 10h, either in DMSO or in 10 μΜ JQ1 and subsequently stained for pH3 by immunofluorescence(n=3 embryos). The Tunel assay was employed to analyse cell death. Apoptotic nuclei were detected in peripheral edges of both DMSO and JQ1 treated explants (n=3 embryos).
B. No significant differences were found between DMSO and JQ1 treated explants in number of pH3 positive loci (explants micro-dissected from 3 different embryos, quantified in 18 optical sections per condition). Data presented as Mean ± SEM.
C. CLE/cPSM explants treated in DMSO (a-d) or with 10 μΜ JQ1 (a’-d’) for 10h analysed by *in situ* hybridisation. At this time frame, both *Sox2* (n=3/3) and *Brachyury* (n=7/10) expression is downregulated, however differentiation markers such as *Pax6* (n=0/5 embryos) and *Paraxis* (n=0/3 embryos) are not expressed in either control DMSO or JQ1 treated explants.
D. CLE/cPSM explants after 10h of culture in DMSO (a-d) or in 10 μΜ JQ1 (a’-b’). Representative *in situ* hybridisation images of explants treated with CT99021 (a,b,a’,b’) or 100 nM RA (c,d,c’,d’) for 14h or 20h post the 10h culture in DMSO or 10 μΜ JQ1.
E. RT-qPCR analysis of gene expression changes in control CLE/cPSM explants cultured in DMSO or in 10 μΜ JQ1 for 10h. Relative gene expression, normalised to actin levels. Data from 3 independent experiments, presented as Mean ± SEM. Scale bars are 100 μm.

The effect of MYC inhibition on cell behaviour in these assays was also addressed. MYC inhibition using JQ1 did not induce apoptosis as revealed by the Tunel assay, with positive cells only detected at the cut edges of explants in both treatment and control conditions (Figure 5A). Myc orchestrates expression of many genes involved in cell cycle progression (Dang et al., 2006, Zeller et al., 2003, Zeller et al., 2006, Eilers and Eisenman, 2008). Analysis of the key known MYC target *cyclin E1* indicated a reduction in transcripts following 10h JQ1 treatment (Figure 5E). We therefore extended this analysis and determining number of pH3 positive cells, indicative of late G2/mitotic phase (Figure 5A). We did not observe significant differences in this time period, however this might be related to the cell cycle length in NMPs, estimated to be approximately 7-8h in the chicken embryo (Olivera-Martinez et al., 2014) and as such it is perhaps not surprising that we do not see proliferation impairment following 10h of MYC activity suppression.

To determine if the lack of precocious differentiation following MYC inhibition was due to a requirement for MYC activity for initiation/progression of differentiation, we first transiently suppressed MYC activity using 10 μΜ JQ1 for 10h, as above, and subsequently washed out the inhibitor and cultured the explants for further 14h either in plain culture medium or in the presence of differentiation stimuli. Wash-out of MYC inhibition was not sufficient to stimulate initiation of expression of differentiation markers (Suppl. Figure 4). To stimulate differentiation towards the mesoderm lineage we employed the potent GSK3 antagonist CT99021 (Cohen and Goedert, 2004) which has been extensively used to generate NMPs (Gouti et al., 2014, Verrier et al., 2017) and PSM (Chal et al., 2015) *in vitro*. We incubated explants (previously treated for 10h with DMSO or JQ1) in 30 μΜ CT99021 for 14h and then checked by ISH for expression of the cPSM marker *Tbx6* (Chapman et al., 1996). DMSO control explants showed high levels of *Tbx6* expression however JQ1 treated explants exhibited very low expression (Figure 5D; a-a’). However, prolonging exposure to CT99021 for a further 6h (20h in total, post-removal of MYC inhibition) induced high *Tbx6* expression in the JQ1 treated explants. In contrast, at this time point the DMSO treated explants no longer expressed *Tbx6*, likely due to their further differentiation along the paraxial mesoderm maturation pathway (Figure 5D; b-b’). We then repeated exactly the same experiment, this time stimulating retinoid signalling, to promote neural differentiation, using 100 nM of Retinoic Acid for 14h and 20h as above. In a similar way, we found that JQ1 treated explants were delayed in their response to upregulate expression of the neural marker gene *Pax6* (Stoykova et al., 1996, Patel et al., 2013) in response to RA stimulation (Figure 5D; c,d,c’d’).

These experiments suggest that Myc directs the expression of multiple genes within the NMP/cPSM network. One of the gene sets involves core factors functioning in NMP (*Fgf8, Wnt3a, Wnt8a, Sox2*) and the cPSM (*Fgf8, Wnt3a*) progenitor pool maintenance. The other gene sets are involved in cell cycle progression (*p21, cyclin E1)* and glycolytic metabolism (*Eno3, Tpi1*). At the same time, MYC activity is a requirement for the differentiation response to external signalling cues, likely by regulating a different target gene set.

### WNT, FGF and NOTCH converge upstream cMyc expression

Since we found that MYC activity is important for the maintenance of the CLE/cPSM signalling network, we then explored what signals regulate cMyc expression in this domains. cMyc has been shown to be a canonical NOTCH and WNT target in many *in vitro* systems of cultured cells (He et al., 1998, Weng et al., 2006, Palomero et al., 2006, Herranz et al., 2014) and FGF/ERK has been propose to act upstream of cMyc transcription (through activation of the downstream ETs factors; (de Nigris et al., 2001), mRNA stability (Notari et al., 2006) and protein turnover (Sears, 2004, Kress et al., 2015). In addition, Retinoid signalling has been previously shown to have a negative effect on Myc expression in teratocarcinoma cells (Griep and DeLuca, 1986). Here we employed gain of function and loss of function approaches to test whether WNT, FGF, NOTCH or RA regulate cMyc transcription in the CLE/cPSM domain.

We first checked whether FGF regulates cMyc by taking explants of CLE/cPSM and either incubating them for 4h with 3 μΜ of the MEK inhibitor PD184352 (Allen et al., 2003) to interfere with FGF signal transduction or with recombinant FGF8 protein to stimulate the endogenous pathway. *Sprouty2* expression, a bona fide FGF/ERK target gene (Sivak et al., Thisse and Thisse, 2005), was monitored in parallel to verify the efficacy of the assays. We find that explants where ERK activity was suppressed showed severe downregulation of both *Sprouty2* expression and *cMyc* expression, whereas explants exposed to exogenous FGF showed increased levels of both genes, indicating a role for FGF acting upstream of *cMyc* (Figure 6 Aa-d’). In a similar manner, we tested whether WNT signalling regulates cMyc expression using 10 μΜ of the known Tankyrase inhibitor XAV939 to suppress WNT signalling (Huang et al., 2009) or WNT3a conditioned medium to stimulate WNT. *Axin2* was used as a readout for WNT activity (Jho et al., 2002). Similarly to the FGF experiment, we found that *Axin2* and *cMyc* were both downregulated following XAV939 treatment and were both upregulated in response to exogenous WNT signalling (Figure 6Ba-d’). We then used the γ-secretase inhibitor LY411575 to inhibit NOTCH signalling (Curry et al., 2005) and found that both *Lfringe* (NOTCH target gene; (McGrew et al., 1998)) and *cMyc* expression was lost in the treated explants. (Figure 6C). Finally, culture of CLE/cPSM explants with 100 nM RA for 6h reduced *Fgf8* levels, as previously reported (Diez del Corral et al., 2003), however it did not confer changes in *cMyc* levels, suggesting that within this progenitors it is unlikely that RA acts upstream cMyc (Figure 6D).

**Figure 6:**
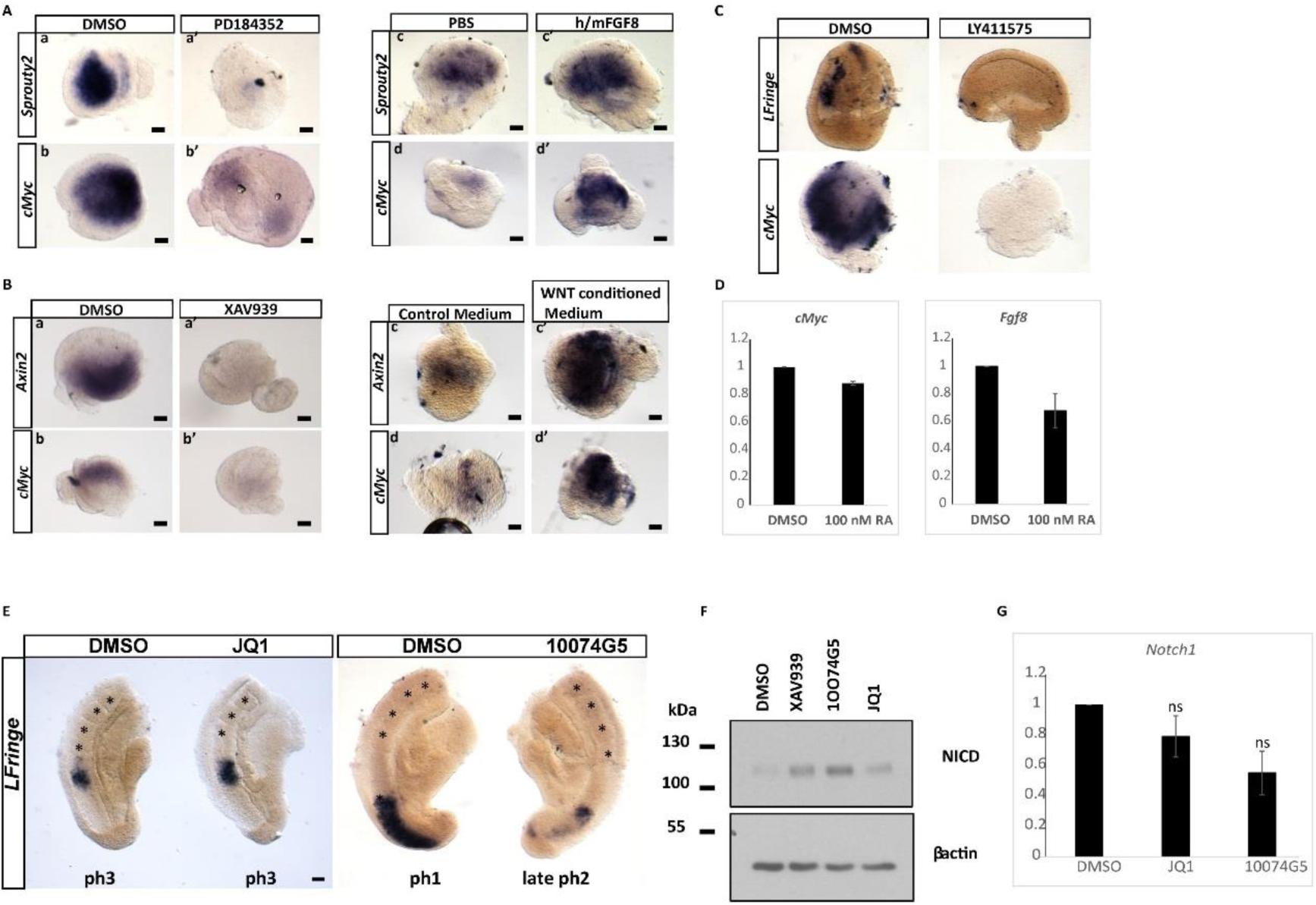
cMyc is co-regulated by WNT, FGF and NOTCH signals and MYC activity suppression results in delays in somite formation. A. Representative *in situ* hybridisation images of CLE/cPSM explants cultured for 4h in DMSO (a,b) or with 3 μΜ PD184352 (a’,b’) to block ERK phosphorylation *Sprouty2* (n=5/5 embryos) and *cMyc* (n=5/6 embryos) expression is downregulated in the PD184352 treated explants. Stimulation of FGF signalling using 250 ng/ml recombinant FGF8 protein for 8h results in upregulation of both *Sprouty2* (n=5/8 embryos) and *cMyc* (n=5/9 embryos) in CLE/cPSM explants (c’,d’) when compared to control explants cultured in PBS (c,d).
B. Representative *in situ* hybridisation images of CLE/cPSM explants cultured for 4h in DMSO (a,b) or 10 μΜ XAV939 (a’,b’) to block Tankyrase activity. *Axin2* (n=7/7 embryos) and *cMyc* (n=4/5 embryos) expression is downregulated in the XAV939 treated explants. Stimulation of WNT signalling using WNT3a conditioned culture medium for 6h results in upregulation of both *Axin2* (n= 3/3embryos) and *cMyc* (n=3/3 embryos) in CLE/cPSM explants (c’,d’) when compared to control explants cultured in DMSO (c,d).
C. Representative *in situ* hybridisation images of E8.5 half-tail explants (micro-dissected at the level below the last formed somite) cultured for 4h in DMSO (a,b) or CLE/cPSM explants cultured in the presence of 250 nM LY411575 (a’,b’) to block γ-secretase activity. *Lfringe* (n=4/4 embryos) and *cMyc* (n=4/4 embryos) expression is downregulated in the LY411575 treated explants compared to controls. (For A,B,C, scale bars are 100 μm)
D. RT-qPCR quantification of *cMyc* and *Fgf8* expression in CLE/cPSM cultured for 6h in DMSO or 100 nM RA. Relative gene expression normalised to actin levels, and presented as Mean ± SEM. Data from 2 independent experiments.
E. Representative images of E9.5 half-tail explants treated with (a) 10 μΜ JQ1 or (b) with 75 μΜ 10074G5 for 4h, labelled by *in situ* hybridisation for the NOTCH clock gene *LFringe*. Myc inhibition delayed *Lfringe* phase pattern expression (11/16 explants treated with JQ1, 4/5 explants treated with 10074G5). Scale bars are 100 μm. Western-Blot analysis for NICD levels on E9.5 embryo tails. Tails that were incubated with JQ1 or 10074G5 for 4h show increased NICD levels when compared to control DMSO tails cultured in parallel.
F. RT-qPCR quantification of *Notch1* expression in control CLE/cPSM explants cultured for 6h in DMSO or in CLE/cPSM explants cultured in parallel in 10 μΜ JQ1 or 75 μΜ 10074G5, reveals no significant expression changes. Relative gene expression normalised to actin levels, and presented as Mean ± SEM. Data from 3 independent experiments.

These experiments suggest that the three segmentation clock pathways (WNT, FGF and NOTCH) converge in regulating *cMyc* transcription in the NMPs and cPSM. However, it is known that there is extensive crosstalk (Bone et al., 2014, Katoh, 2007, Stulberg et al., 2012, Akai et al., 2005), so each of these pathways could be acting directly or indirectly to regulate cMyc expression.

### MYC inhibition leads to delayed clock gene oscillations and slows somitogenesis

We then asked whether MYC activity plays a role in the process of somite formation. To test this, we bissected E9.5 embryo tails and incubated them for 4h with one half cultured in the presence of DMSO and the other half cultured in the presence of the 10 μΜ JQ1 or 75 μΜ 10074G5. We then processed the half tail explants for *in situ* hybridisation and looked at the phase expression patterns of the NOTCH target clock gene *Lfringe*. The somite number in each explant was counted before and after culture. Control explants had formed a least one new somite during the 4h incubation, however, explants where MYC activity was inhibited displayed delayed oscillations of *Lfringe* expression as compared to their control counterparts and in some cases also formed less somites (Figure 6E).

We have previously published that tight regulation of NICD (NOTCH1 intracellular domain) turnover regulates the period of clock gene oscillations (Wiedermann et al., 2015). To see if the observed delay in somite formation and clock gene oscillations upon MYC inhibition is linked to NOTCH signalling we analysed levels of NICD in E9.5 tails cultured for 4h in the presence or absence of MYC inhibitors by Western blotting. We found that tail explant lysates incubated with either of the 2 small molecule inhibitors displayed higher levels of NICD, when compared to DMSO treated tail lysates (Figure 6F). In addition, NICD levels between JQ1 or 10074G5 treated tail lysates were similar to NICD levels of tail explant lysates treated with XAV939 (Figure 6F), a factor previously shown to affect the clock pace through increasing NICD levels/stability by reducing high NICD turnover (Wiedermann et al., 2015). Previously cMyc has been shown to repress *Notch1* transcription (Zinin et al., 2014). To test whether the observed increases in NICD levels upon MYC activity suppression result from increases in *Notch1* transcription, we analysed the levels of *Notch1* in CLE/cPSM explants treated for 6h with 10 μΜ JQ1 or 75 μΜ 10075G5. We did not find significant changes in *Notch1* transcript levels upon MYC activity suppression (Figure 6G). These data suggest that the increases in NICD levels following MYC activity suppression are due to posttranslational effects are not a result of increased levels of *Notch1* transcription.

### Conditional inducible cMyc depletion results in reduction of *Fgf8* expression levels

To further study these diverse functions *in vivo*, we generated a conditional inducible mouse line in which we could specifically genetically ablate cMyc expression in a spatially and temporally controlled manner from the tail of the post-implantation embryo. To this end we employed an available transgenic cMyc mouse line, where lox-P sites have been placed on either side of the exons two and three of the cMyc locus (Trumpp et al. (2001); hereafter called cMyc^FL/FL^ allele). To specifically ablate cMyc from the embryonic tail, we used an available mouse line, whereby CRE expression is under the control of the Nkx1.2 promoter (Rodrigo Albors et al., 2016). Nkx1.2 starts to be expressed at the primitive streak and posterior epiblast at E7, and is expressed in the entire CLE and in the pre-Neural Tube domain at E8.5 (Rodrigo Albors et al., 2016). Induction of homologous recombination through the CRE recombinase activity results in excision of the two exons and loss of function of the cMyc genetic product, (Trumpp et al., 2001). Visualization of CRE activation domain was possible through immunofluorescence labelling for the YFP protein; the YFP allele, controlled by the Rosa26 promoter, was activated for expression upon CRE mediated excision of a lox-P flanked STOP sequence located upstream the YFP locus (Rodrigo Albors et al., 2016).

Using *in situ* hybridization on E8.5 embryos we verified a complete loss of *cMyc* transcript in the CLE of cMyc KO embryos, whereas gene expression was present in the somites and head (Supl. Figure 6A). In addition, immunofluorescence labelling for GFP revealed intense staining in the CLE domain, with a few cells labelled in the neural tube in accordance to Rodrigo Albors et al. (2016). The cMyc KO embryos were indistinguishable from control wild type embryos and loss of cMyc did not affect formation of PSM, somites and neural tissue. These experiments suggest that acute cMyc loss from the E8.5 CLE does not impact short term embryo development.

In order to investigate whether loss of cMyc from the E8.5 tail region might impact embryo development during body axis elongation stages we collected embryos on E9.5, E10.5 and E11.5 dpc. We found that cMyc KO embryos were of normal morphology, and formed all CLE derivatives (PSM, somites and neural tissue) normally (Figure 7Aa, Ba’).

**Figure 7:**
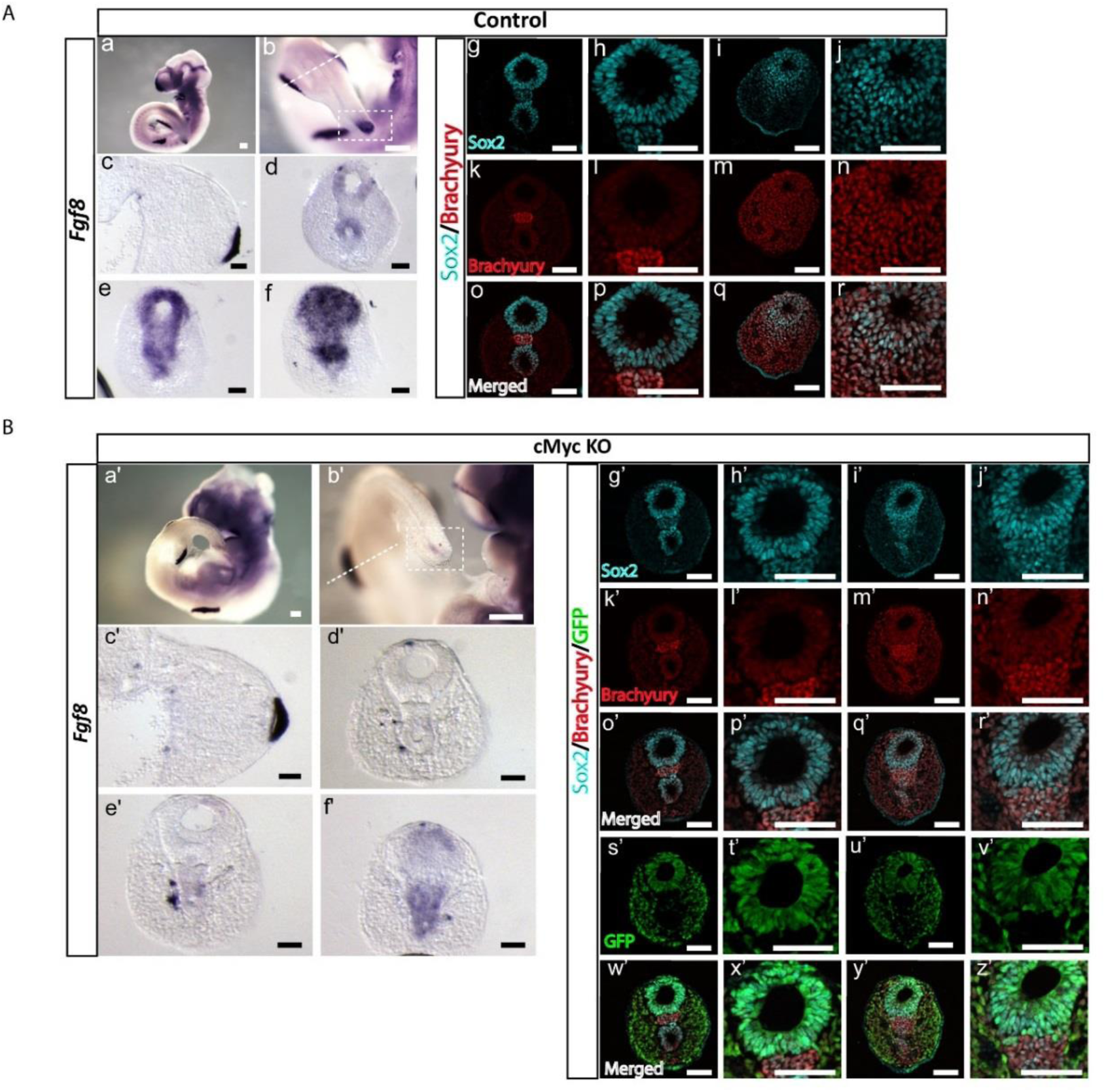
Conditional inducible cMyc depletion from the CLE results in downregulation of *Fgf8* expression during axis elongation. A. (a-f) Representative images of E10.5 Control embryos (n=3 embryos) labelled by *in situ* hybridization for *Fgf8* expression show high expression levels in the hind limbs and in the tailbud. (c) is a transverse section in the limb bud region demarcated by the white dotted line in (b). (d-f) are transverse sections of the CNH and tailbud demarcated by white dotted box in (b). (g-r) Are confocal images on transverse sections of the CNH and tailbud labelled by immunofluorescence for Sox2 and Brachyury and show large number of co-expressing cells in the tailbud mesenchyme (data from 10 confocal sections from 2 embryos).
B. Representative images of E10.5 cMyc conditional inducible mutant embryos (n=3 embryos) labelled by *in situ* hybridization for *Fgf8* expression (a’-f’) show high levels in the hind limbs, whereas very low levels are detected in the tailbud. (c’) is a transverse section in the limb bud region demarcated by the white dotted line in (b’). (d’-f’) are transverse sections of the CNH and tailbud demarcated by white dotted box in (b’). (g’-z’) Confocal images on transverse sections of the CNH and tailbud labelled by immunofluorescence for Sox2, Brachyury and GFP show large number of co-expressing cells in the tailbud mesenchyme. GFP cells are mostly absent from the domain of the CNH (s’), which coincides with the domain where *Fgf8* expression is still detected (f’) (data from 10 confocal sections from 2 embryos). Scale bars are 100 μm.

Strikingly we found a profound reduction of *Fgf8* expression in the PSM of the E9.5 cMyc KO embryos (n=5 embryos; Supl. Figure 6B). In particular, *Fgf8* expression was severely downregulated in the overlying caudal most neuroepithelium whereas the PSM was completely devoid of transcript expression. We then checked E10.5 cMyc embryos, which were of normal morphology, and similarly to E9.5 embryos, they express severely reduced *Fgf8* levels compared to control embryos where *in situ* hybridization colour revelation was conducted in parallel (n=3 embryos) (Figure 7Ab-e, Bb’-e’). *Fgf8* expression was spared only in the mesodermal compartment of the CNH (Figure 7Bf’). *Fgf8* expression in the limb buds of E10.5 cMyc KO embryos was unaffected as compared to control embryos (Figure 7Aa-c, Ba’-c’).

In order to address whether cMyc depletion and subsequent *Fgf8* downregulation affected the presence or number of Sox2/Brachyury co-expressing NMP cells, we carried out immunofluorescence labelling in E10.5 tails for Sox2, Brachyury and GFP. We found that GFP expressing cells in the tailbud (indicative of cMyc depletion) largely co-expressed Sox2 and Brachyury, in a similar manner to wild type embryos, suggesting that cMyc mediated loss of *Fgf8* throughout almost all of the tail end of the embryo is insufficient to directly affect the NMP identity (Fig 7Ag-r, Bg’-z’). Interestingly, the mesenchymal domain that still expresses low levels of *Fgf8* in the cMyc-KO embryos (Figure 7Bf’), is almost devoid of GFP-expressing cells (Figure 7Bs’), in accordance with Rodrigo Albors et al. (2016) who show that Nkx1.2 expressing cells do not contribute to this mesoderm compartment of the CNH. These FGF8 positive cells may serve in part to maintain the NMP progenitors in the KO embryos. To further decipher whether other NMP regulatory factors could compensate for *Fgf8* loss, we checked the expression of *Wnt3a* in KO embryos. Expression of this gene appeared unaffected (Suppl. Figure 6) suggesting that further compensatory mechanisms safeguard expression of this gene. It is possible that the prolonged persistence of the *Fgf8* mRNA (Dubrulle and Pourquie, 2004) which is still expressed in low levels in the cMyc-KO embryos could indirectly promote *Wnt3a* expression as extensive crosstalk between WNT and FGF signals has previously been reported in these tissues (reviewed n Aulehla and Pourquié (2010), Henrique et al. (2015), Gibb et al. (2010)).

## Discussion

cMyc has been reported to be expressed in the early post-implantation epiblast (Claveria et al., 2013, Sancho et al., 2013) and to be a pluripotency factor in embryonic stem cells (Chappell and Dalton, 2013). Interestingly the E8.5 CLE is a region where the pluripotency factor Oct4 is still expressed, whereas its expression is downregulated in the next developmental stage (E9.5) (Aires et al., 2016). Thus, cMyc expression in the CLE coincides with the presence of other known pluripotency factors (Sox2, Oct4), further highlighting that the CLE shares characteristics of the early pluripotent epiblast (Henrique et al., 2015, Gouti et al., 2015). However, in contrast to Oct4 (Aires et al., 2016), we found that cMyc continues to be expressed in the tailbud during tail growth stages from E9.5 onwards. Using pharmacological strategies to inhibit MYC activity (Yin et al., 2003, Delmore et al., 2011), we found that MYC functions to promote expression of factors that maintain the NMP pool (by sustaining *Wnt3a/8a* and *Fgf8* expression) and the NMP identity (Sox2/ Brachyury co-expression) by promoting *Sox2* expression. Interestingly, cMyc has been shown to positively regulate Sox2 expression (Lin et al., 2009) and to directly stimulate expression of WNT components while suppressing transcription of WNT inhibitors (Fagnocchi et al., 2016) in mouse pluripotent stem cells. In the same context, cMyc was found to be WNT regulated, and it in turn was shown to establish a positive feedback WNT network that was indispensable for ESC identity maintenance (Fagnocchi and Zippo, 2017, Fagnocchi et al., 2016). Similarly, in our work we find that both WNT and FGF converge upstream of cMyc transcription suggesting that cMyc could be establishing and maintaining a Myc/WNT/FGF network operating in the NMPs. Given that secreted WNT/FGF act on the underlying mesoderm and control PSM maturation, it could be possible that the aforementioned network operates in both of these adjacent progenitor domains (NMPs/cPSM), even though active transcription (of at least a subset of FGF components) is restricted to the overlying CLE (Dubrulle and Pourquie, 2004). In addition to WNT and FGF, we found that NOTCH activity is also important for cMyc transcription, in alignment with other published work identifying cMyc as a canonical NOTCH target, where again it operates in a positive feedforward loop (Palomero et al., 2006).

Very recently, two independent studies have highlighted a novel role for glycolytic metabolism, during axis elongation in chicken and mouse embryos (Bulusu et al., 2017, Oginuma et al., 2017). The authors found that during axis elongation, a gradient of glycolytic activity, downstream FGF signaling, is established across the PSM, with higher metabolic rates in the tailbud and decreasing levels towards the anterior of the PSM. Inhibition of glycolysis resulted in loss of NMPs, premature differentiation towards the neural lineage and cessation of elongation (Oginuma et al., 2017). MYC has been shown to be involved in the regulation of metabolism in other cell contexts (Claveria et al., 2013, Dang et al., 2006, Eilers and Eisenman, 2008, Fagnocchi and Zippo, 2017, Hsieh et al., 2015) while recently cMyc was found to link FGF signaling and glycolysis during vascular development in mice (Yu et al., 2017). In our study we found that MYC activity is crucial for maintenance of mRNA levels of 2 key glycolytic genes (Tpi1, Eno3) that have been shown to exhibit graded rostro-caudal expression along the PSM (Oginuma et al., 2017). Taken together these data suggest that, in the NMP and caudal PSM populations, MYC activity may regulate progenitor pool maintenance through the integration of proliferative WNT/FGF signals, as well as regulation of a specific subset of metabolic genes.

A striking finding was that MYC activity suppression resulted in delays of dynamic segmentation clock gene expression across the PSM, which also coincided with increases in NICD levels. Interestingly, MYC activity suppression did not result in significant changes in *Notch1* mRNA levels, in contrast to a previous study indicating cMyc is a negative regulator of *Notch1* expression in the chicken embryo neural tube (Zinin et al., 2014). A possible explanation for how MYC might affect NICD levels in the PSM could be through negative regulation of proteins mediating NICD turnover. We have previously shown that WNT or CDK inhibition (Gibb et al., 2009, Wiedermann et al., 2015) delays the periodicity of dynamic *LFringe* clock gene expression across the PSM and that this phenotype is linked to inefficient NICD turnover. Given we demonstrate here that cMyc expression is WNT regulated in this tissue and we find that MYC regulates transcription of the CDK inhibitor p21 it is possible to hypothesise that MYC acts downstream of WNT signaling and upstream of p21, which in turn controls downstream CDKs that regulate NICD phosphorylation and subsequent turnover. It would be interesting to investigate further whether MYC activity is crucial for proper timing of FGF and WNT “clock” gene oscillations, which would provide insight into how the segmentation clock pathways are mechanistically linked.

Since cMyc is highly expressed in the CLE and persists in the tailbud (in contrast to *MycN*), we chose to acutely deplete cMyc from the embryonic tail region and study possible defects caused through cMyc loss during the body axis elongation stages. Global cMyc KO embryos, are smaller than wild-type embryos and display multiple defects including abnormalities in neural tube closure (Davis et al., 1993). They do however survive up to E10.5, possibly due to compensatory activity of MycN, which shows increased expression in response to cMyc depletion (Trumpp et al., 2001). Thus, it is possible that our conditional inducible cMyc KO embryos lack a very profound phenotype due to MycN activity which is still operational. We were however able to highlight a direct requirement for cMyc, as a transcriptional regulator of *Fgf8* in the caudal neuroepithelium and tailbud in regions where the Sox2/Brachyury co-expressing NMPS reside. *Fgf8* expression was spared only in the mesodermal compartment of the CNH of KO embryos. Despite *Fgf8* loss throughout most of this caudal domain, the NMP pool was not compromised (at least at E10.5), which could be attributed either to compensation from the low level of FGF8 remaining, or other FGF factors that are present in the embryonic tail, or to uninterrupted WNT activity (for example through *Wnt3a*), which was not affected following genetic depletion of cMyc in this region and has been shown to be crucial for the NMP pool maintenance (Garriock et al., 2015). In addition, Fgf8 has been previously shown to be dispensable for body axis elongation in the mouse (Perantoni et al., 2005) even though it required for initial body axis specification and mesoderm migration through the primitive streak (Sun et al., 1999).

In summary, the current work uncovered multiple novel functional roles for MYC activity during embryonic body axis elongation. Future work should focus on deciphering the global Myc transcriptional signature within the progenitors that mediate this process.

## Materials and Methods

### Mouse Embryo Collection and explant dissection

Pregnant female mice were culled by cervical dislocation and the uteri were dissected and collected in Phosphate Buffer Saline (PBS). Embryos of either E8.5, E9.5, E10.5 or E11.5 days post-coitum (dpc) were washed in Leibovitz’s L15 medium (Life Technologies) or DMEM-F12 + 0.1 % Glutamax medium (Life Technologies) and collected in L15 or DMEM-F12 supplemented with 5-10% Foetal Bovine Serum (Life Technologies. All animal procedures were carried out under the Animal Scientific procedures Act (1987).

For E8.5 CLE/cPSM explant dissection mouse embryos were collected in L15 medium (Life Technologies). The embryos were first staged according to the somite number and embryos at the 5-7 somite stage were used. The tissue flanking the node and rostral to the tailbud mesoderm which included the caudal lateral epiblast and underlying mesoderm was dissected and subsequently bisected along the longitudinal axis of the embryo to give right and left caudal embryo explant pairs. The explants were then individually embedded in freshly prepared rat-tail derived collagen mix (consisting of 71.5% rat tail collagen (Corning), 23.8% 5xL15, 23.8% Acetic Acid, 4.7% NaHCO3) and cultured for the appropriate timeframe at 37 °C in a controlled atmosphere of 5% CO_2_ in air in a culture medium consisting of Optimem (Life Technologies) supplemented with 5% FBS, 2 μΜ glutamine and 50 μΜ/ml gentamycin. For each pair, one explant was cultured in the appropriate volume of DMSO diluted in the above culture medium or in culture medium supplemented with the appropriate dilution of a small molecule inhibitor to suppress MYC, WNT, NOTCH or FGF/ERK activity, described in the Table 1. For WNT signalling stimulation, culture medium derived from L Cells (ATCC^®^ CRL-2648™; hereafter called Control medium) or in DMEM based WNT3a enriched medium derived from L Wnt-3A cells (ATCC® CRL-2647™, hereafter called WNT conditioned medium), supplemented with 10% FBS (Labtech), 2 mM L-glutamine (Lonza,) 1% penicillin/streptomycin (Lonza) and was prepared in house and used on CLE/cPSM explants. For FGF stimulation experiments, 250 ng/ml were employed as described in Diez del Corral et al. (2003).

**Table 1:**
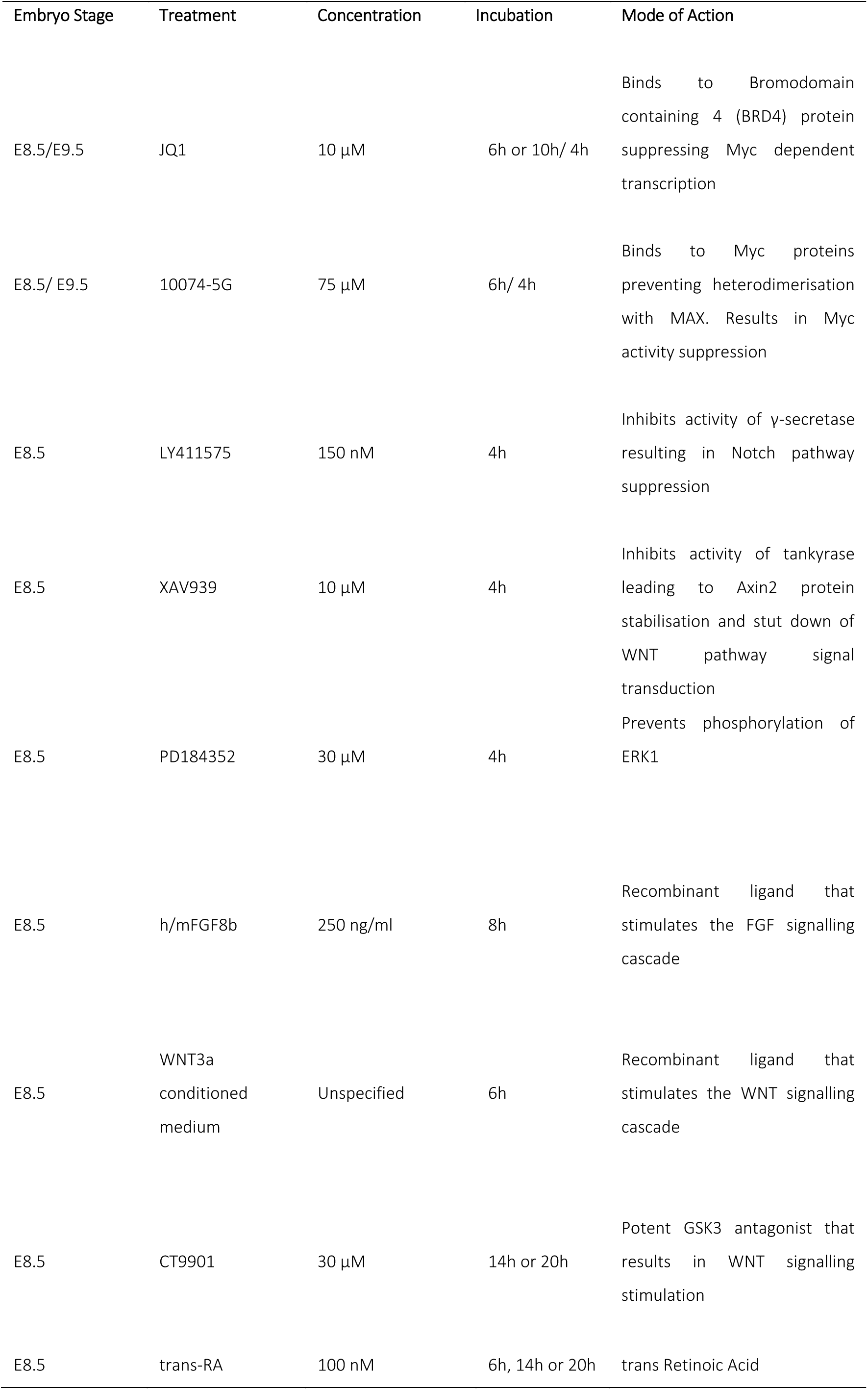
Summary of embryo explant treatments

At the end of the culture time, the embedded explants were fixed in 4% Formaldehyde (Sigma) or 4% Paraformaldehyde (Electron Microscopy) diluted in PBS, either for 2 hours at room temperature or overnight at 4 °C, if they were to be processed for *in situ* hybridisation or for 2 hours at 4°C if they were to be processed for immunofluorescence. They were directly lysed in RLT Buffer (Qiagen) if they were to be processed for RT-qPCR analysis. E9.5 half tail explants, were micro-dissected and cultured as described in Bone et al. (2014).

### Differentiation of CLE/caudal PSM explants from E8.5 embryos towards mesoderm or neural lineages

CLE/cPSM explants derived from E8.5 embryos were cultured for 10h at 37 °C as described above. At the end of the 10h incubation, both control (DMSO) and JQ1 treated collagen embedded explants were rinsed in clean PSB and washed in PBs twice for 5 min. Subsequently, fresh culture medium (Optimem; Life Technologies); 5% FBS, 2 μΜ glutamine and 50 μΜ/ml gentamycin) supplemented either with 30 μΜ CT9901 (Tocris) or with 100 nM RA (Sigma) was added to the explants and they were further cultured for 14h or 20 hours at 37 °C in an controlled atmosphere of 5% CO_2_ in air.

At the end of the culture time, the explants were fixed in 4% Formaldehyde (Sigma) or 4% Paraformaldehyde (Electron Microscopy) diluted in PBS, overnight at 4 °C, and were processed for *in situ* hybridisation.

### *In situ* Hybridisation

For *in situ* hybridization (ISH), the embryos and embryo derived explants were first fixed overnight in 4% PFA at 4 °C. ISH was carried out following standard procedures. Colour revelation was performed after an overnight incubation with 1/1000 dilution of the anti-digoxigenin antibody conjugated to alkaline phosphatase (Anti-Digoxigenin-AP, Fab fragments, Promega). Control and inhibitor treated explants or wild type and genetically modified embryos were always treated in parallel and colour revelation was initiated and stopped simultaneously.

### Immunocytochemistry

E8.5 wholemount, cryosections or explants of embryos were stained for immunofluorescence using primary antibodies against Sox2 (neural marker; raised in goat, Immune Systems), Brachyury (mesoderm marker; raised in rabbit, Santa Cruz biotechnology or raised in goat, R&D Systems) and Phospo Histone 3 (pH3; Upstate). cMyc positive cells were labelled using the polyclonal anti c-Myc antibody (raised in rabbit, Abcam). Briefly, embryos and explants were blocked overnight in 10% Heat inactivated Donkey serum diluted in PBT (1% Tween, 1% Triton, 2% Bovine serum Albumin diluted in PBS) at 4 °C. All primary antibodies were employed in a final concentration 1/200 in blocking solution and the embryos or explants were incubated overnight at 4°C. Following washes with PBST, the anti-donkey fluorescently conjugated secondary antibodies Alexa 488 and Alexa 594 (Invitrogen) were added in a final dilution 1/200 in blocking solution and the tissues were incubated overnight at 4 °C. Nuclei were counterstained using Dapi in a final 1/1000 dilution.

### Tunel Assay

The tunel assay was performed following a standard protocol optimized for tissue processing and using the ApopTag Kit (Millipore) according manufacturers instructions.

### Culture and Differentiation of H9 Human Pluripotent stem cells (PSC) to the NMP state

H9 hESC were maintained as feeder-free cultures in DEF-based medium (Cellartis DEF-CS) supplemented with 30 ng/ml bFGF (Peprotech) and Noggin (10ng/ml, Peprotech) on fibronectin coated plates, and enzymatically passaged using TryPLselect (Thermofisher), and differentiated to the NMP state following the protocol from Verrier et al. (2017).

### Image acquisition and analysis

Images of fluorescently labelled wholemount, sections and explants of E8.5 embryos were taken using a Zeiss 710 confocal microscope equipped with a LASOS camera. Images of ISH were acquired using the Leitz DM RB Leica microscope, which is equipped with a Nicon D1X camera. Figure preparation was carried out using the free online software OMERO (www.openmicroscopy.org).

### Quantitative Real-Time Polymerase Chain Reaction (RT-qPCR)

Total RNA from H9 Human Pluripotent stem cells and human NMPs or derived explants (6xCLE/cPSM explants per sample) was purified using the Qiagen miniKit (Qiagen) following manufacturers instructions and concentration and quality of the eluted RNA was determined using the NANOdrop System. cDNA synthesis was performed using the Superscript III kit (Invitrogen) in 0.25 ug, 0.5 ug or 1 ug of purified RNA according to manufacturers instructions. qRTPCR was performed using 1ul of synthesised cDNA and 9 ul of Power SYBR® Green PCR Master Mix (Life Technologies/ Fisher) diluted in which were primers against the gene of interest (Table 2). A CFX96 thermocycler (Bio Rad) was used for target cDNA amplification. Dilution curves for each primer set were performed to ensure that the primers were working in 100% efficiency and the del-delcT (Livak and Schmittgen, 2001) method was used to analyse gene expression levels. Statistical significance was assessed either with the student T-test of with Analysis of Variance (ANOVA).

**Table 2:**
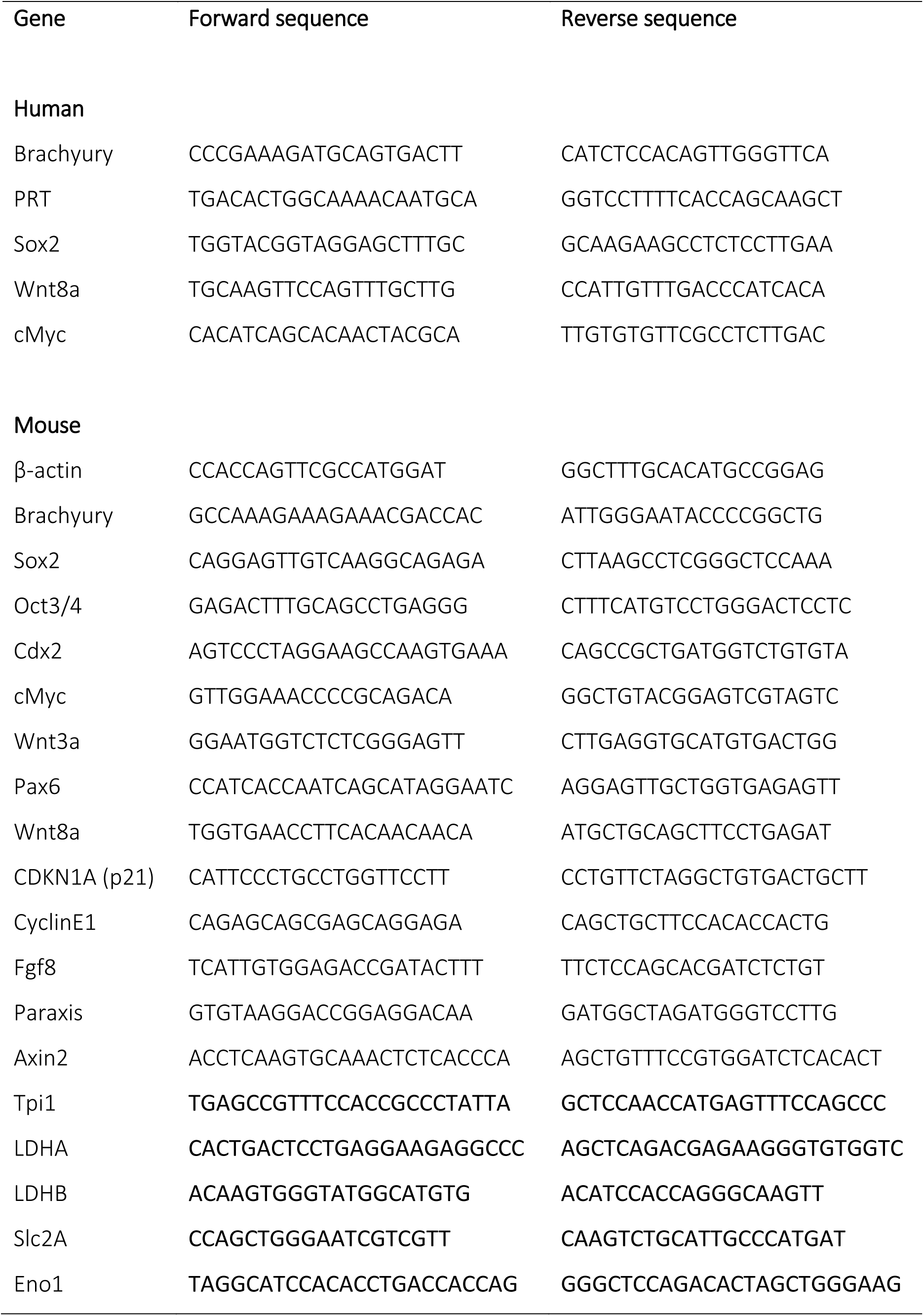
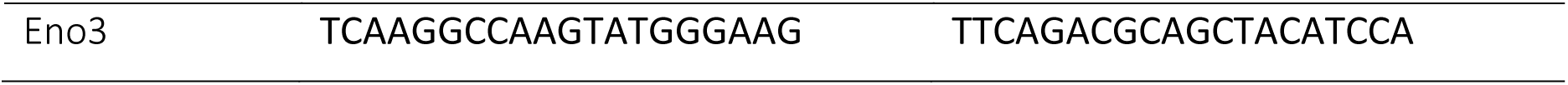
List of Primers used

### Western Blotting of mouse E9.5 embryo tails

Protein extraction and Western Blotting was carried out following standard procedures using 10 μg of protein and the following dilutions of antibodies: 1:1000 of the rabbit α-mouse NICD (Cell Signalling), and 1:10000 mouse α-βactin (Proteintech).

### Generation of the inducible conditional cMyc mouse line

In order to genetically deplete cMyc from the tail end of the embryo, we utilised two available C56 mouse lines. One previously generated in the lab (Rodrigo Albors et al., 2016) where CRE recombinase expression is driven under the control of the Nkx 1.2 promoter (hereafter called Nkx1.2 ^ERT2_CRE^ locus) which also carried LoxP-flanked STOP sequence upstream of a EYFP reporter transgene of the Rosa26 ubiquitous promoter (hereafter called YFP locus) (Srinivas et al., 2001, Rodrigo Albors et al., 2016) and one where LoxP sites have been placed on either site of the exon2 and exon3 of the cMyc locus (hereafter called cMyc FLOX^N^ locus)(Trumpp et al., 2001). Homozygote mice for the cMyc FLOX^N^, Nkx1.2 ^ERT2_CRE^, YFP alleles were morphologically normal and indistinguishable from C57 wild type mice. Induction of homologous recombination was achieved by administering 200 ul of Tamoxifen (Sigma) diluted in 10% ETOH/ 90% vegetable oil in a final concentration of 40 ug/ml to the pregnant mother on E6 and E7 dpc by oral gavage. Embryos were collected and analysed on E8.5, E9.5, E10.5 and E11.5 dpc. Validation of the region were the cMyc locus was knocked-out was achieved by immunofluorescence labelling against the YFP protein and by ISH against *cMyc*. Throughout the generation and maintenance of the mouse colony, ear biopsies were dissected from mouse pups for genotyping. DNA was extracted from the ear biopsies by direct addition of 20 ul of the MICROLYSIS mix (Cambio) directly in the biopsy sample. The PCR programmes and primer sets for detection of the cMyc FLOX^N^ locus are from Trumpp et al. (2001) and for the Nkx1.2 ^ERT2_CRE^ and YFP alleles are from Rodrigo Albors et al. (2016).

## Acknowledgements

We are grateful to all members of the Dale and Storey Laboratories for input and discussions. We are especially thankful to P. Halley for cloning cMyc; to F.A Carrierri for assistance with Western blotting; to P. Bozatzi for helping with the WNT3a conditioned medium; to R. Elliot and N. Suttie for assistance with ISH.

## Competing Interests

The authors declare no competing interests

## Funding

This work was supported by a Cancer Research PhD studentship to I.M; by a Wellcome Trust Strategic award [097945/Z/11/Z] to J.K.D; and by a Wellcome Trust Investigator Award to KGS [WT102817AIA].

## Suppl. Figure Legends

**Sup Figure 1.**
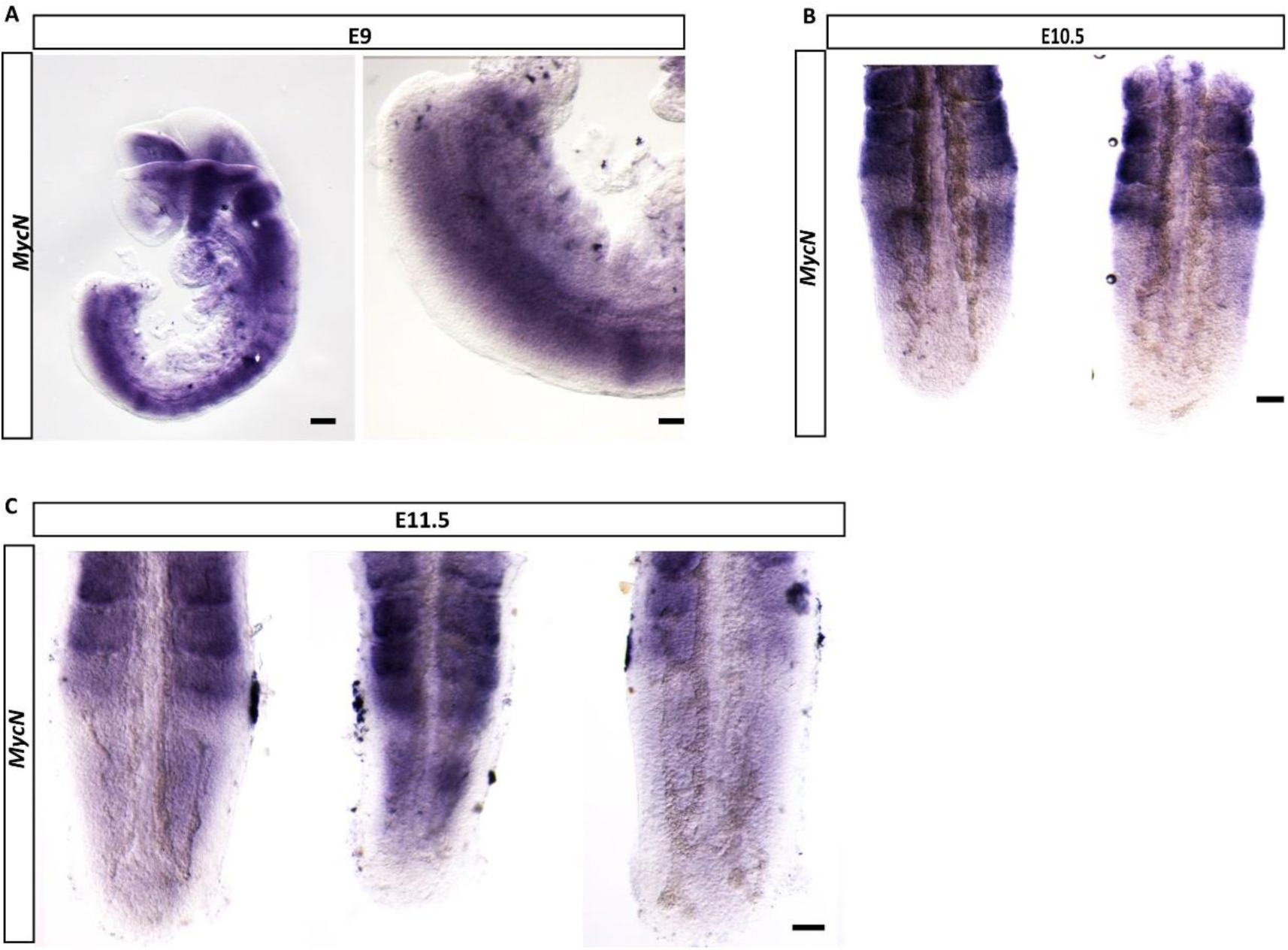
MycN expression at E9-E11.5. A. Representative *in situ* hybridisation images for *MycN* at E9.5 (n=3) reveal an enrichment for *MycN* expression in the PSM compartment, and in particular a strong rostral domain of expression below the level of the last somite (white arrowhead), similarly to cMyc.
B. Representative *in situ* hybridisation images of different embryo tails at E10.5 (n=5) *MycN* is highly expressed in the somites of the E10.5 embryo tails and in variable levels across the PSM. A strong band of mRNA expression was evident in some tails, below the level of the last somite (white arrowhead).
C. Representative *in situ* hybridisation images of different embryo tails at E11.5 (n=4) days of development. *MycN* is highly expressed in the somites of the E11.5 embryo tails and in variable levels across the PSM. A strong band of mRNA expression was evident in some tails, below the level of the last somite (white arrowhead).

**Sup Figure 2.**
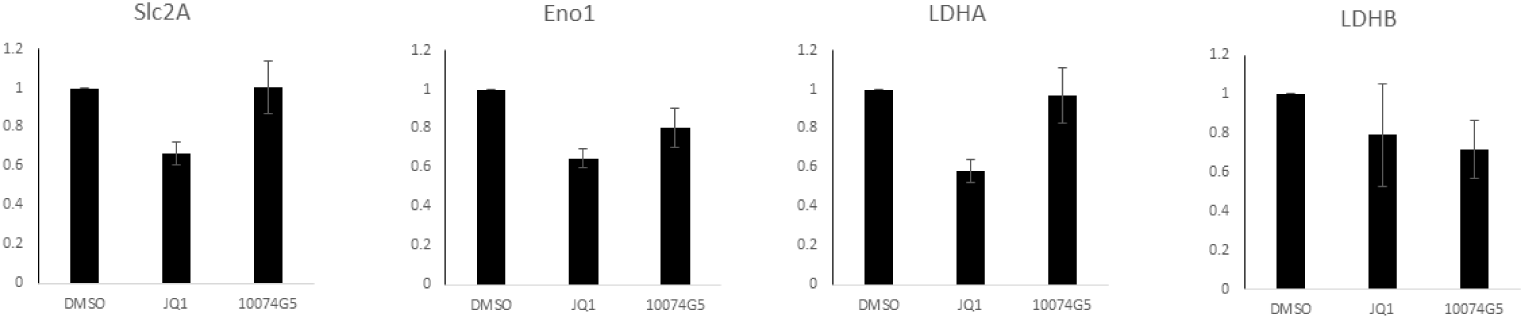
Expression levels of metabolic genes that are insensitive to 6h MYC activity suppression. Relative expression normalised to actin levels, data from 3 independent experiments, presented as Mean ± SEM.

**Sup Fig 3.**
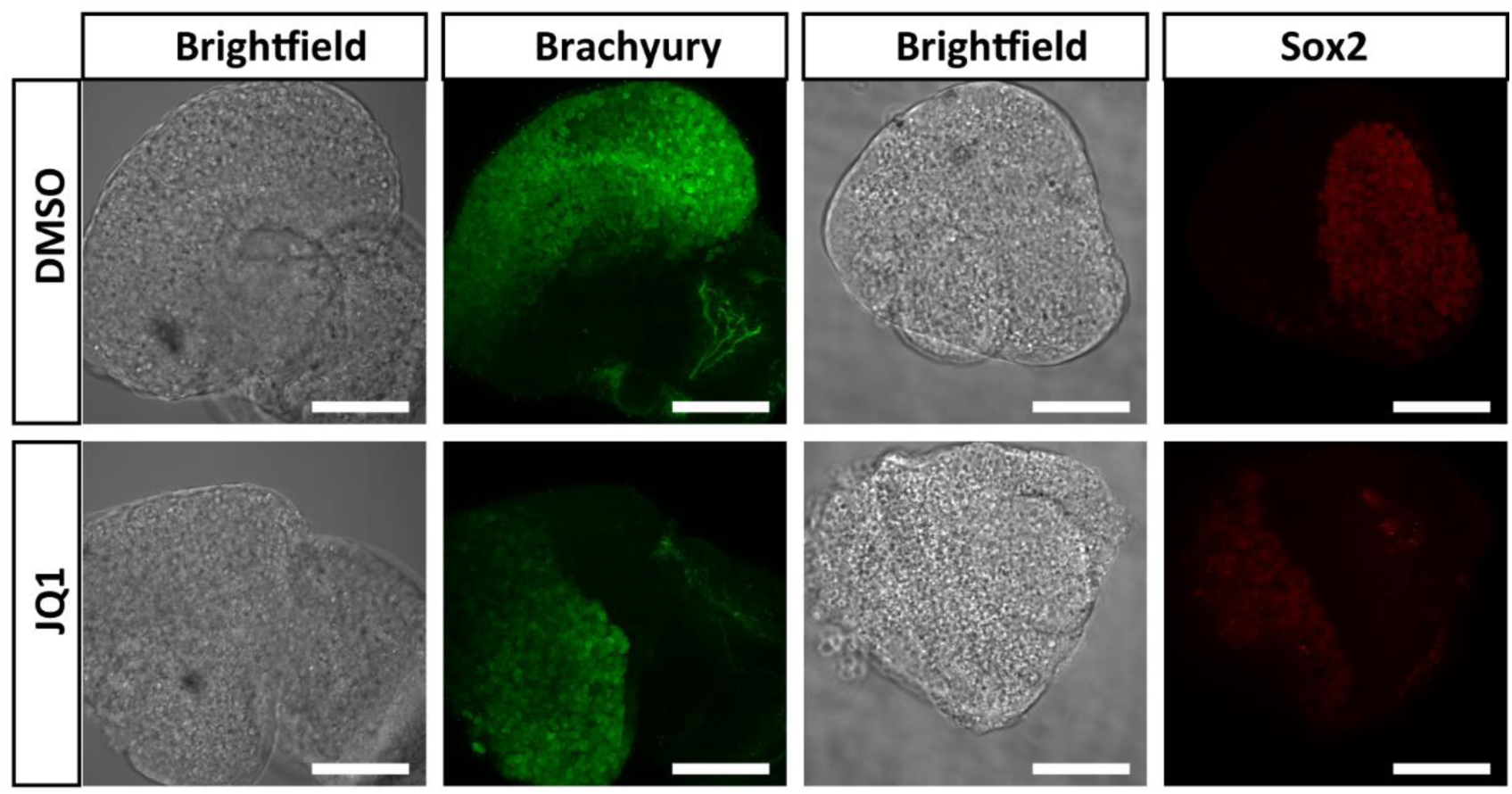
Sox2 and Brachyury expression at 10h of JQ1 mediated MYC activity suppression. Representative confocal images of CLE/cPSM explant pairs treated for 10h with 10 μΜ JQ1 or DMSO show that, Brachyury protein levels are similar between control and JQ1 treated explants (n=3/4 embryos). Sox2 protein levels are mildly downregulated in JQ1 treated explants (n=6/6 embryos). Scale bars are 100 μm.

**Sup Figure 4.**
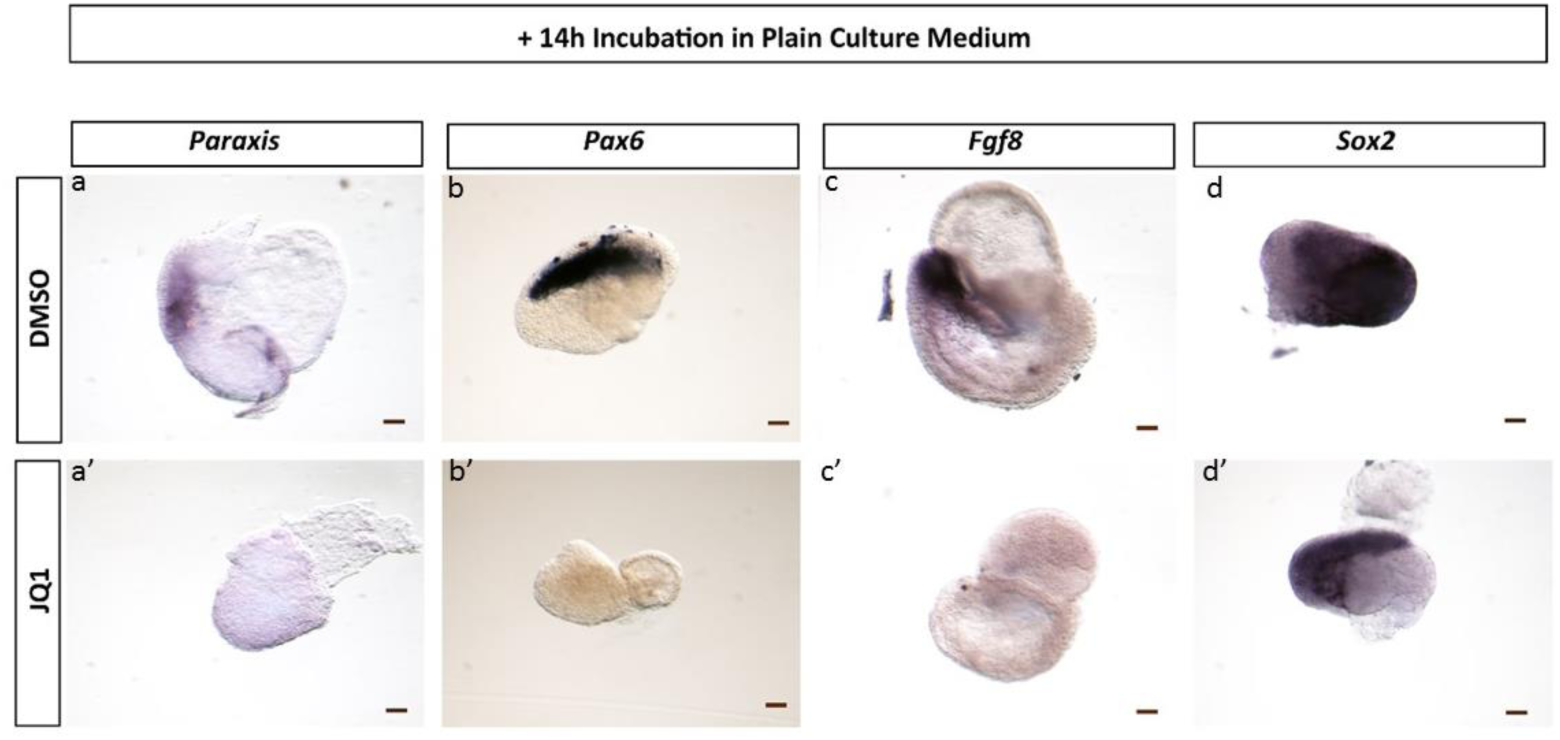
Wash out of JQ1 does not promote differentiation. Representative *in situ* hybridisation images of CLE/cPSM explants that were treated with (<u>a’,b’,c’,d’</u>) 10 μΜ JQ1 or (<u>a,b,c,d</u>) equivalent DMSO volume for 10h and subsequently cultured for further 14h in plain culture medium. In that timeframe explants that were initially incubated with JQ1 do not upregulate expression of *Paraxis* (n=0/2 embryos) or *Pax6* (n=0/3 embryos). Furthermore, they do not recover *Fgf8* expression (n=0/7 embryos) but they do express *Sox2* (n=4/4 embryos) at similar levels to their control counterparts. At this timeframe, JQ1 treated explants appear to be of a smaller size than the control DMSO explants. Scale bars are 100 μm.

**Sup Figure 5:**
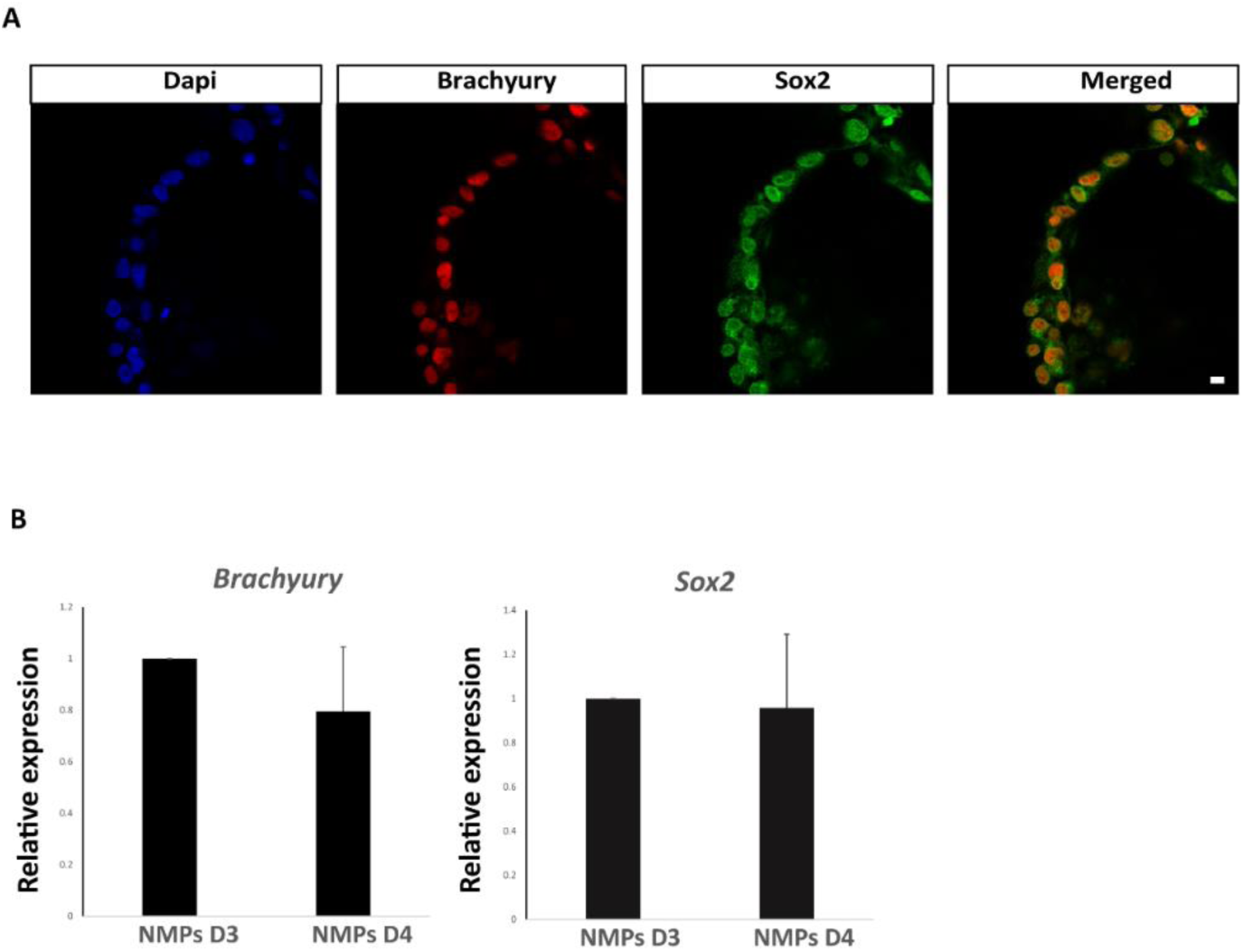
hNMP maintenance for 24h. A. Representative confocal images of *in vitro* generated hNMPs from H9 pluripotent stem cells, show co-localisation of Sox2/Brachyury positive cells on Day3 of the differentiation protocol. Scale bar is 100 μm.
B. qRTPCR expression analysis of Sox2 and Brachyury between Day3 and Day4 of the differentiation protocol shows that gene expression levels for these two genes remain unaltered in the 24h timeframe (data from two independent experiments, expression normalised to levels of PRT2 gene)

**Sup Figure 6:**
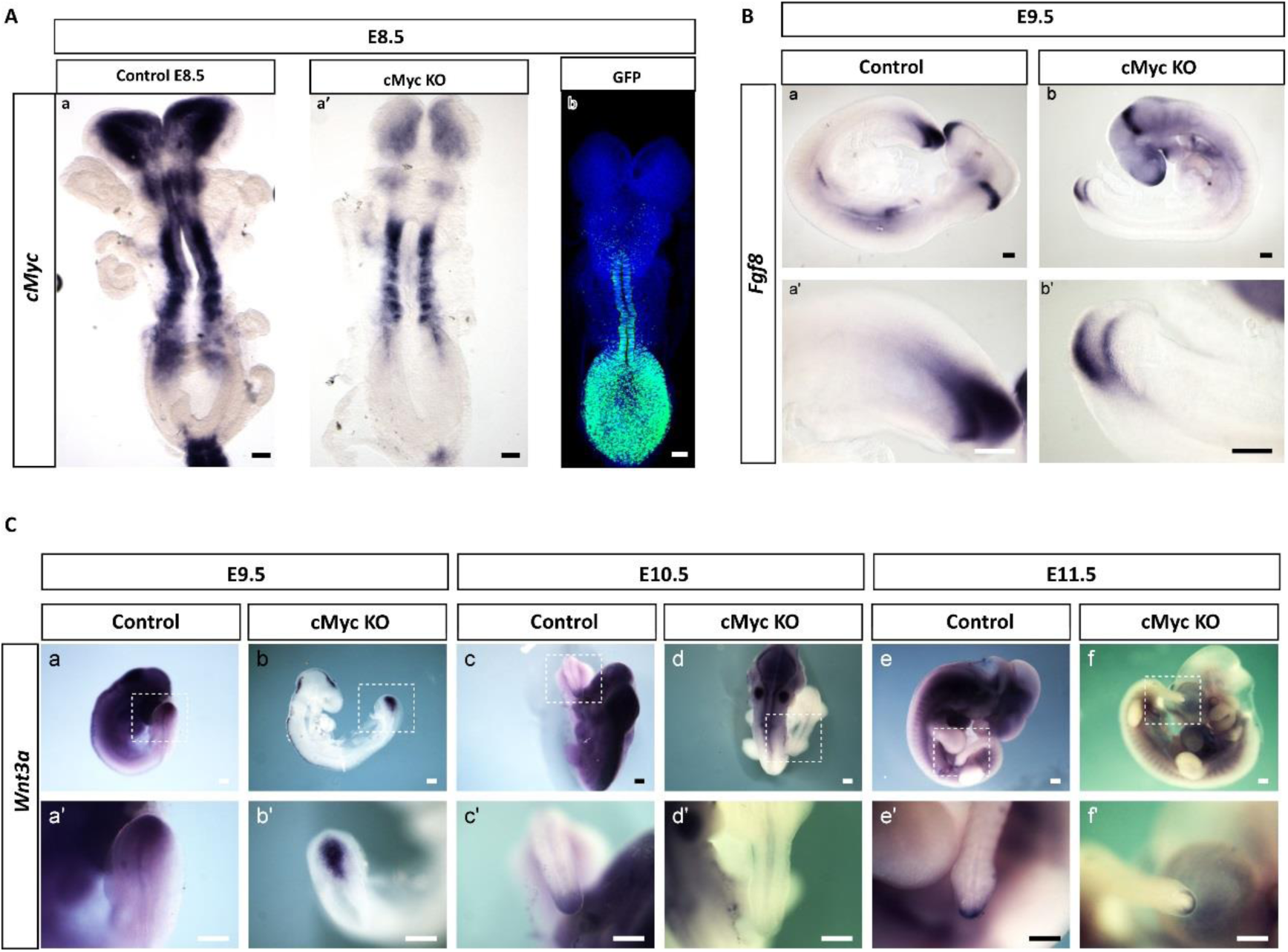
*Fgf8* but not *Wnt3a* is downregulated in conditional inducible cMyc KO embryos. A. Representative *in situ* hybridisation images of (<u>a</u>) Control Wild type (n=5) and (<u>b</u>) cMyc KO embryos (n=5) where a reduction of *cMyc* expression in the tail region is evident. Tamoxifen was administered at E6.5 and E7.5 days of development. (b) Representative staining by immunofluorescence for GFP labels the CLE cells where recombination and subsequent cMyc depletion has taken place (n=5 embryos).
B. Representative *in situ* hybridisation images for *Fgf8* expression in control (<u>a,a’</u>) and cMyc KO embryos (<u>b,b’</u>) at E9.5 show a reduction in *Fgf8* transcript levels specifically in the tail region (n=4 control and n=5 cMyc KO embryos)
C. Representative images of control (<u>a,c,e</u>) and cMyc KO (<u>b,d,f</u>) E9.5, E10.5 and E11.5 embryos labelled by *in situ* hybridisation for *Wnt3a* expression (<u>a’-f’</u> panels are higher magnification images of the area denominated with the white dotted rectangle on <u>a-f</u>). We find very mild downregulation of *Wnt3a* expression in 1 out of 2 E9.5 embryos, 2/5 E10.5 and no downregulation in 3/3 E11.5 embryos. Scale bars are 100 μm.

## References

Aires, R., Jurberg, A. D., Leal, F., Nóvoa, A., Cohn, M. J. & Mallo, M. 2016. Oct4 Is a Key Regulator of Vertebrate Trunk Length Diversity. Developmental Cell, 38, 262–274.

Akai, J., Halley, P. A. & Storey, K. G. 2005. FGF-dependent Notch signaling maintains the spinal cord stem zone. Genes & Development, 19, 2877–2887.

Alitalo, K., Bishop, J. M., Smith, D. H., Chen, E. Y., Colby, W. W. & Levinson, A. D. 1983. Nucleotide sequence to the v-myc oncogene of avian retrovirus MC29. Proc Natl Acad Sci U S A, 80, 100–4.

Allen, L. F., Sebolt-Leopold, J. & Meyer, M. B. 2003. CI-1040 (PD184352), a targeted signal transduction inhibitor of MEK (MAPKK). Seminars in Oncology, 30, 105–116.

Aulehla, A. & Pourquié, O. 2010. Signaling Gradients during Paraxial Mesoderm Development. Cold Spring Harbor Perspectives in Biology, 2, a000869.

Blackwood, E. M. & Eisenman, R. N. 1991. Max: a helix-loop-helix zipper protein that forms a sequence-specific DNA-binding complex with Myc. Science, 251, 1211–7.

Blackwood, E. M., Luscher, B., Kretzner, L. & Eisenman, R. N. 1991. The Myc:Max protein complex and cell growth regulation. Cold Spring Harb Symp Quant Biol, 56, 109–17.

Bone, R. A., Bailey, C. S. L., Wiedermann, G., Ferjentsik, Z., Appleton, P. L., Murray, P. J., Maroto, M. & Dale, J. K. 2014. Spatiotemporal oscillations of Notch1, Dll1 and NICD are coordinated across the mouse PSM. Development (Cambridge, England), 141, 4806–4816.

Brodeur, G. M., Seeger, R. C., Schwab, M., Varmus, H. E. & Bishop, J. M. 1984. Amplification of N-myc in untreated human neuroblastomas correlates with advanced disease stage. Science, 224, 1121–4.

Bulusu, V., Prior, N., Snaebjornsson, M. T., Kuehne, A., Sonnen, K. F., Kress, J., Stein, F., Schultz, C., Sauer, U. & Aulehla, A. 2017. Spatiotemporal Analysis of a Glycolytic Activity Gradient Linked to Mouse Embryo Mesoderm Development. Developmental Cell, 40, 331–341.e4.

Burgess, R., Cserjesi, P., Ligon, K. L. & Olson, E. N. 1995. Paraxis: A Basic Helix-Loop-Helix Protein Expressed in Paraxial Mesoderm and Developing Somites. Developmental Biology, 168, 296–306.

Cambray, N. & Wilson, V. 2002. Axial progenitors with extensive potency are localised to the mouse chordoneural hinge. Development, 129.

Cambray, N. & Wilson, V. 2007. Two distinct sources for a population of maturing axial progenitors. Development, 134, 2829.

Chal, J., Oginuma, M., Al Tanoury, Z., Gobert, B., Sumara, O., Hick, A., Bousson, F., Zidouni, Y., Mursch, C., Moncuquet, P., Tassy, O., Vincent, S., Miyanari, A., Bera, A., Garnier, J.-M., Guevara, G., Hestin, M., Kennedy, L., Hayashi, S., Drayton, B., Cherrier, T., Gayraud-Morel, B., Gussoni, E., Relaix, F., Tajbakhsh, S. & Pourquie, O. 2015. Differentiation of pluripotent stem cells to muscle fiber to model Duchenne muscular dystrophy. Nat Biotech, 33, 962–969.

Chapman, D. L., Agulnik, I., Hancock, S., Silver, L. M. & Papaioannou, V. E. 1996. Tbx6, a mouse T-Box gene implicated in paraxial mesoderm formation at gastrulation. Developmental biology, 180, 534–542.

Chappell, J. & Dalton, S. 2013. Roles for MYC in the establishment and maintenance of pluripotency. Cold Spring Harb Perspect Med, 3, a014381.

Claassen, G. F. & Hann, S. R. 2000. A role for transcriptional repression of p21(CIP1) by c-Myc in overcoming transforming growth factor β-induced cell-cycle arrest. Proceedings of the National Academy of Sciences of the United States of America, 97, 9498–9503.

Claveria, C., Giovinazzo, G., Sierra, R. & Torres, M. 2013. Myc-driven endogenous cell competition in the early mammalian embryo. Nature, 500, 39–44.

Cohen, P. & Goedert, M. 2004. GSK3 inhibitors: development and therapeutic potential. Nat Rev Drug Discov, 3, 479–487.

Curry, C. L., Reed, L. L., Golde, T. E., Miele, L., Nickoloff, B. J. & Foreman, K. E. 2005. Gamma secretase inhibitor blocks Notch activation and induces apoptosis in Kaposi’s sarcoma tumor cells. Oncogene, 24, 6333–6344.

Dale, J. K., Maroto, M., Dequeant, M. L., Malapert, P., Mcgrew, M. & Pourquie, O. 2003. Periodic notch inhibition by lunatic fringe underlies the chick segmentation clock. Nature, 421, 275–8.

Dang, C. V., O’Donnell, K. A., Zeller, K. I., Nguyen, T., Osthus, R. C. & Li, F. 2006. The c-Myc target gene network. Semin Cancer Biol, 16, 253–64.

Davis, A. C., Wims, M., Spotts, G. D., Hann, S. R. & Bradley, A. 1993. A null c-myc mutation causes lethality before 10.5 days of gestation in homozygotes and reduced fertility in heterozygous female mice. Genes Dev, 7, 671–82.

De Nigris, F., Mega, T., Berger, N., Barone, M. V., Santoro, M., Viglietto, G., Verde, P. & Fusco, A. 2001. Induction of ETS-1 and ETS-2 Transcription Factors Is Required for Thyroid Cell Transformation. Cancer Research, 61, 2267.

Delfino-Machín, M., Lunn, J. S., Breitkreuz, D. N., Akai, J. & Storey, K. G. 2005. Specification and maintenance of the spinal cord stem zone. Development, 132, 4273.

Delmore, J. E., Issa, G. C., Lemieux, M. E., Rahl, P. B., Shi, J., Jacobs, H. M., Kastritis, E., Gilpatrick, T., Paranal, R. M., Qi, J., Chesi, M., Schinzel, A. C., Mckeown, M. R., Heffernan, T. P., Vakoc, C. R., Bergsagel, P. L., Ghobrial, I. M., Richardson, P. G., Young, R. A., Hahn, W. C., Anderson, K. C., Kung, A. L., Bradner, J. E. & Mitsiades, C. S. 2011. BET bromodomain inhibition as a therapeutic strategy to target c-Myc. Cell, 146, 904–17.

Dequeant, M. L., Glynn, E., Gaudenz, K., Wahl, M., Chen, J., Mushegian, A. & Pourquie, O. 2006. A complex oscillating network of signaling genes underlies the mouse segmentation clock. Science, 314, 1595–8.

Dequeant, M. L. & Pourquie, O. 2008. Segmental patterning of the vertebrate embryonic axis. Nat Rev Genet, 9.

Deschamps, J. & Duboule, D. 2017. Embryonic timing, axial stem cells, chromatin dynamics, and the Hox clock. Genes & Development, 31, 1406–1416.

Diez Del Corral, R., Olivera-Martinez, I., Goriely, A., Gale, E., Maden, M. & Storey, K. 2003. Opposing FGF and retinoid pathways control ventral neural pattern, neuronal differentiation, and segmentation during body axis extension. Neuron, 40.

Downs, K. M., Martin, G. R. & Bishop, J. M. 1989. Contrasting patterns of myc and N-myc expression during gastrulation of the mouse embryo. Genes Dev, 3, 860–9.

Dubrulle, J. & Pourquie, O. 2004. fgf8 mRNA decay establishes a gradient that couples axial elongation to patterning in the vertebrate embryo. Nature, 427, 419–422.

Eilers, M. & Eisenman, R. N. 2008. Myc's broad reach. Genes Dev, 22, 2755–66.

Emanuel, B. S., Balaban, G., Boyd, J. P., Grossman, A., Negishi, M., Parmiter, A. & Glick, M. C. 1985. N-myc amplification in multiple homogeneously staining regions in two human neuroblastomas. Proc Natl Acad Sci U S A, 82, 3736–40.

Fagnocchi, L., Cherubini, A., Hatsuda, H., Fasciani, A., Mazzoleni, S., Poli, V., Berno, V., Rossi, R. L., Reinbold, R., Endele, M., Schroeder, T., Rocchigiani, M., Szkarlat, Z., Oliviero, S., Dalton, S. & Zippo, A. 2016. A Myc-driven self-reinforcing regulatory network maintains mouse embryonic stem cell identity. Nat Commun, 7, 11903.

Fagnocchi, L. & Zippo, A. 2017. Multiple Roles of MYC in Integrating Regulatory Networks of Pluripotent Stem Cells. Front Cell Dev Biol, 5, 7.

Garriock, R. J., Chalamalasetty, R. B., Kennedy, M. W., Canizales, L. C., Lewandoski, M. & Yamaguchi, T. P. 2015. Lineage tracing of neuromesodermal progenitors reveals novel Wnt-dependent roles in trunk progenitor cell maintenance and differentiation. Development (Cambridge, England), 142, 1628–1638.

Gartel, A. L., Ye, X., Goufman, E., Shianov, P., Hay, N., Najmabadi, F. & Tyner, A. L. 2001. Myc represses the p21((WAF1/CIP1)) promoter and interacts with Sp1/Sp3. Proceedings of the National Academy of Sciences of the United States of America, 98, 4510–4515.

Gibb, S., Maroto, M. & Dale, J. K. 2010. The segmentation clock mechanism moves up a notch. Trends in Cell Biology, 20, 593–600.

Gibb, S., Zagorska, A., Melton, K., Tenin, G., Vacca, I., Trainor, P., Maroto, M. & Dale, J. K. 2009. Interfering with Wnt signalling alters the periodicity of the segmentation clock. Developmental Biology, 330, 21–31.

Goto, H., Kimmey, S. C., Row, R. H., Matus, D. Q. & Martin, B. L. 2017. FGF and canonical Wnt signaling cooperate to induce paraxial mesoderm from tailbud neuromesodermal progenitors through regulation of a two-step epithelial to mesenchymal transition. Development, 144, 1412.

Gouti, M., Delile, J., Stamataki, D., Wymeersch, F. J., Huang, Y., Kleinjung, J., Wilson, V. & Briscoe, J. 2017. A Gene Regulatory Network Balances Neural and Mesoderm Specification during Vertebrate Trunk Development. Dev Cell, 41, 243–261 e7.

Gouti, M., Metzis, V. & Briscoe, J. 2015. The route to spinal cord cell types: a tale of signals and switches. Trends in Genetics, 31, 282–289.

Gouti, M., Tsakiridis, A., Wymeersch, F. J., Huang, Y., Kleinjung, J., Wilson, V. & Briscoe, J. 2014. In Vitro Generation of Neuromesodermal Progenitors Reveals Distinct Roles for Wnt Signalling in the Specification of Spinal Cord and Paraxial Mesoderm Identity. PLOS Biology, 12, e1001937.

Griep, A. E. & Deluca, H. F. 1986. Decreased c-myc expression is an early event in retinoic acid-induced differentiation of F9 teratocarcinoma cells. Proceedings of the National Academy of Sciences of the United States of America, 83, 5539–5543.

He, T.-C., Sparks, A. B., Rago, C., Hermeking, H., Zawel, L., Da Costa, L. T., Morin, P. J., Vogelstein, B. & Kinzler, K. W. 1998. Identification of c-<em>MYC</em> as a Target of the APC Pathway. Science, 281, 1509–1512.

Henrique, D., Abranches, E., Verrier, L. & Storey, K. G. 2015. Neuromesodermal progenitors and the making of the spinal cord. Development, 142, 2864–75.

Herranz, D., Ambesi-Impiombato, A., Palomero, T., Schnell, S. A., Belver, L., Wendorff, A. A., Xu, L., Castillo-Martin, M., Llobet-Navas, D., Cordon-Cardo, C., Clappier, E., Soulier, J. & Ferrando, A. A. 2014. A NOTCH1-driven MYC enhancer promotes T cell development, transformation and acute lymphoblastic leukemia. Nat Med, 20, 1130–1137.

Horne, G. A., Stewart, H. J. S., Dickson, J., Knapp, S., Ramsahoye, B. & Chevassut, T. 2014. Nanog Requires BRD4 to Maintain Murine Embryonic Stem Cell Pluripotency and Is Suppressed by Bromodomain Inhibitor JQ1 Together with Lefty1. Stem Cells and Development, 24, 879–891.

Hsieh, A. L., Walton, Z. E., Altman, B. J., Stine, Z. E. & Dang, C. V. 2015. MYC and metabolism on the path to cancer. Semin Cell Dev Biol, 43, 11–21.

Huang, S.-M. A., Mishina, Y. M., Liu, S., Cheung, A., Stegmeier, F., Michaud, G. A., Charlat, O., Wiellette, E., Zhang, Y., Wiessner, S., Hild, M., Shi, X., Wilson, C. J., Mickanin, C., Myer, V., Fazal, A., Tomlinson, R., Serluca, F., Shao, W., Cheng, H., Shultz, M., Rau, C., Schirle, M., Schlegl, J., Ghidelli, S., Fawell, S., Lu, C., Curtis, D., Kirschner, M. W., Lengauer, C., Finan, P. M., Tallarico, J. A., Bouwmeester, T., Porter, J. A., Bauer, A. & Cong, F. 2009. Tankyrase inhibition stabilizes axin and antagonizes Wnt signalling. Nature, 461, 614–620.

Hubaud, A. & Pourquie, O. 2014. Signalling dynamics in vertebrate segmentation. Nat Rev Mol Cell Biol, 15, 709–21.

Ikegaki, N., Minna, J. & Kennett, R. H. 1989. The human L-myc gene is expressed as two forms of protein in small cell lung carcinoma cell lines: detection by monoclonal antibodies specific to two myc homology box sequences. EMBO J, 8, 1793–9.

Jho, E.-H., Zhang, T., Domon, C., Joo, C.-K., Freund, J.-N. & Costantini, F. 2002. Wnt/β-Catenin/Tcf Signaling Induces the Transcription of Axin2, a Negative Regulator of the Signaling Pathway. Molecular and Cellular Biology, 22, 1172–1183.

Kato, K., Kanamori, A., Wakamatsu, Y., Sawai, S. & Kondoh, H. 1991. Tissue Distribution of N-myc Expression in the Early Organogenesis Period of the Mouse Embryo. Development, Growth & Differentiation, 33, 29–39.

Katoh, M. 2007. Networking of WNT, FGF, Notch, BMP, and Hedgehog Signaling Pathways during Carcinogenesis. Stem Cell Reviews, 3, 30–38.

Kerosuo, L. & Bronner, M. E. 2016. cMyc Regulates the Size of the Premigratory Neural Crest Stem Cell Pool. Cell Rep, 17, 2648–2659.

Kim, J.-W., Zeller, K. I., Wang, Y., Jegga, A. G., Aronow, B. J., O’Donnell, K. A. & Dang, C. V. 2004. Evaluation of Myc E-Box Phylogenetic Footprints in Glycolytic Genes by Chromatin Immunoprecipitation Assays. Molecular and Cellular Biology, 24, 5923–5936.

Kress, T. R., Sabo, A. & Amati, B. 2015. MYC: connecting selective transcriptional control to global RNA production. Nat Rev Cancer, 15, 593–607.

Krol, A. J., Roellig, D., Dequeant, M. L., Tassy, O., Glynn, E., Hattem, G., Mushegian, A., Oates, A. C. & Pourquie, O. 2011. Evolutionary plasticity of segmentation clock networks. Development, 138, 2783–92.

Lin, C.-H., Lin, C., Tanaka, H., Fero, M. L. & Eisenman, R. N. 2009. Gene Regulation and Epigenetic Remodeling in Murine Embryonic Stem Cells by c-Myc. PLOS ONE, 4, e7839.

Livak, K. J. & Schmittgen, T. D. 2001. Analysis of relative gene expression data using real-time quantitative PCR and the 2(-Delta Delta C(T)) Method. Methods, 25, 402–8.

Ma, M., Zhao, K., Wu, W., Sun, R. & Fei, J. 2014. Dynamic expression of N-myc in mouse embryonic development using an enhanced green fluorescent protein reporter gene in the N-myc locus. Dev Growth Differ, 56, 152–60.

Maroto, M., Bone, R. A. & Dale, J. K. 2012. Somitogenesis. Development, 139, 2453–6.

Mcgrew, M. J., Dale, J. K., Fraboulet, S. & Pourquie, O. 1998. The lunatic fringe gene is a target of the molecular clock linked to somite segmentation in avian embryos. Curr Biol, 8, 979–82.

Meyer, N. & Penn, L. Z. 2008. Reflecting on 25 years with MYC. Nat Rev Cancer, 8, 976–90.

Naiche, L. A., Holder, N. & Lewandoski, M. 2011. FGF4 and FGF8 comprise the wavefront activity that controls somitogenesis. Proceedings of the National Academy of Sciences, 108, 4018–4023.

Nau, M. M., Brooks, B. J., Battey, J., Sausville, E., Gazdar, A. F., Kirsch, I. R., Mcbride, O. W., Bertness, V., Hollis, G. F. & Minna, J. D. 1985. L-myc, a new myc-related gene amplified and expressed in human small cell lung cancer. Nature, 318, 69–73.

Notari, M., Neviani, P., Santhanam, R., Blaser, B. W., Chang, J.-S., Galietta, A., Willis, A. E., Roy, D. C., Caligiuri, M. A., Marcucci, G. & Perrotti, D. 2006. A MAPK/HNRPK pathway controls BCR/ABL oncogenic potential by regulating MYC mRNA translation. Blood, 107, 2507–2516.

Oginuma, M., Moncuquet, P., Xiong, F., Karoly, E., Chal, J., Guevorkian, K. & Pourquié, O. 2017. A Gradient of Glycolytic Activity Coordinates FGF and Wnt Signaling during Elongation of the Body Axis in Amniote Embryos. Developmental Cell, 40, 342–353.e10.

Olivera-Martinez, I., Harada, H., Halley, P. A. & Storey, K. G. 2012. Loss of FGF-Dependent Mesoderm Identity and Rise of Endogenous Retinoid Signalling Determine Cessation of Body Axis Elongation. PLOS Biology, 10, e1001415.

Olivera-Martinez, I., Schurch, N., Li, R. A., Song, J., Halley, P. A., Das, R. M., Burt, D. W., Barton, G. J. & Storey, K. G. 2014. Major transcriptome re-organisation and abrupt changes in signalling, cell cycle and chromatin regulation at neural differentiation <em>in vivo</em>. Development.

Olivera-Martinez, I. & Storey, K. G. 2007. Wnt signals provide a timing mechanism for the FGF-retinoid differentiation switch during vertebrate body axis extension. Development, 134, 2125–2135.

Palomero, T., Lim, W. K., Odom, D. T., Sulis, M. L., Real, P. J., Margolin, A., Barnes, K. C., O’Neil, J., Neuberg, D., Weng, A. P., Aster, J. C., Sigaux, F., Soulier, J., Look, A. T., Young, R. A., Califano, A. & Ferrando, A. A. 2006. NOTCH1 directly regulates c-MYC and activates a feed-forward-loop transcriptional network promoting leukemic cell growth. Proc Natl Acad Sci U S A, 103, 18261–6.

Patel, N. S., Rhinn, M., Semprich, C. I., Halley, P. A., Dollé, P., Bickmore, W. A. & Storey, K. G. 2013. FGF Signalling Regulates Chromatin Organisation during Neural Differentiation via Mechanisms that Can Be Uncoupled from Transcription. PLOS Genetics, 9, e1003614.

Perantoni, A. O., Timofeeva, O., Naillat, F., Richman, C., Pajni-Underwood, S., Wilson, C., Vainio, S., Dove, L. F. & Lewandoski, M. 2005. Inactivation of FGF8 in early mesoderm reveals an essential role in kidney development. Development, 132, 3859–71.

Perez-Roger, I., Solomon, D. L., Sewing, A. & Land, H. 1997. Myc activation of cyclin E/Cdk2 kinase involves induction of cyclin E gene transcription and inhibition of p27(Kip1) binding to newly formed complexes. Oncogene, 14, 2373–81.

Posternak, V. & Cole, M. D. 2016. Strategically targeting MYC in cancer. F1000Research, 5, F1000 Faculty Rev–408.

Rodrigo Albors, A., Halley, P. A. & Storey, K. G. 2016. Fate mapping caudal lateral epiblast reveals continuous contribution to neural and mesodermal lineages and the origin of secondary neural tube. bioRxiv.

Sakai, Y., Meno, C., Fujii, H., Nishino, J., Shiratori, H., Saijoh, Y., Rossant, J. & Hamada, H. 2001. The retinoic acid-inactivating enzyme CYP26 is essential for establishing an uneven distribution of retinoic acid along the anterio-posterior axis within the mouse embryo. Genes Dev, 15.

Sancho, M., Di-Gregorio, A., George, N., Pozzi, S., Sanchez, J. M., Pernaute, B. & Rodriguez, T. A. 2013. Competitive interactions eliminate unfit embryonic stem cells at the onset of differentiation. Dev Cell, 26, 19–30.

Sawai, S., Shimono, A., Wakamatsu, Y., Palmes, C., Hanaoka, K. & Kondoh, H. 1993. Defects of embryonic organogenesis resulting from targeted disruption of the N-myc gene in the mouse. Development, 117, 1445–55.

Scognamiglio, R., Cabezas-Wallscheid, N., Thier, Marc C., Altamura, S., Reyes, A., Prendergast, áine M., Baumgärtner, D., Carnevalli, Larissa S., Atzberger, A., Haas, S., Von Paleske, L., Boroviak, T., Wörsdörfer, P., Essers, Marieke A. G., Kloz, U., Eisenman, Robert N., Edenhofer, F., Bertone, P., Huber, W., Van Der Hoeven, F., Smith, A. & Trumpp, A. 2016. Myc Depletion Induces a Pluripotent Dormant State Mimicking Diapause. Cell, 164, 668–680.

Sears, R. C. 2004. The Life Cycle of C-Myc: From Synthesis to Degradation. Cell Cycle, 3, 1131–1135.

Sivak, J. M., Petersen, L. F. & Amaya, E. FGF Signal Interpretation Is Directed by Sprouty and Spred Proteins during Mesoderm Formation. Developmental Cell, 8, 689–701.

Srinivas, S., Watanabe, T., Lin, C.-S., William, C. M., Tanabe, Y., Jessell, T. M. & Costantini, F. 2001. Cre reporter strains produced by targeted insertion of EYFP and ECFP into the ROSA26 locus. BMC Developmental Biology, 1, 4.

Staller, P., Peukert, K., Kiermaier, A., Seoane, J., Lukas, J., Karsunky, H., Moroy, T., Bartek, J., Massague, J., Hanel, F. & Eilers, M. 2001. Repression of p15INK4b expression by Myc through association with Miz-1. Nat Cell Biol, 3, 392–9.

Stine, Z. E., Walton, Z. E., Altman, B. J., Hsieh, A. L. & Dang, C. V. 2015. MYC, Metabolism, and Cancer. Cancer discovery, 5, 1024–1039.

Stoykova, A., Fritsch, R., Walther, C. & Gruss, P. 1996. Forebrain patterning defects in Small eye mutant mice. Development, 122, 3453.

Stulberg, M. J., Lin, A., Zhao, H. & Holley, S. A. 2012. Crosstalk between Fgf and Wnt signaling in the zebrafish tailbud. Developmental biology, 369, 298–307.

Sun, X., Meyers, E. N., Lewandoski, M. & Martin, G. R. 1999. Targeted disruption of Fgf8 causes failure of cell migration in the gastrulating mouse embryo. Genes & Development, 13, 1834–1846.

Takahashi, K. & Yamanaka, S. 2006. Induction of pluripotent stem cells from mouse embryonic and adult fibroblast cultures by defined factors. Cell, 126, 663–76.

Tansey, W. P. 2014. Mammalian MYC Proteins and Cancer. New Journal of Science, 2014, 27.

Thisse, B. & Thisse, C. 2005. Functions and regulations of fibroblast growth factor signaling during embryonic development. Dev Biol, 287, 390–402.

Trumpp, A., Refaeli, Y., Oskarsson, T., Gasser, S., Murphy, M., Martin, G. R. & Bishop, J. M. 2001. c-Myc regulates mammalian body size by controlling cell number but not cell size. Nature, 414, 768–73.

Tsakiridis, A., Huang, Y., Blin, G., Skylaki, S., Wymeersch, F., Osorno, R., Economou, C., Karagianni, E., Zhao, S., Lowell, S. & Wilson, V. 2014. Distinct Wnt-driven primitive streak-like populations reflect in vivo lineage precursors. Development, 141, 1209–21.

Turner, D. A., Hayward, P. C., Baillie-Johnson, P., Rué, P., Broome, R., Faunes, F. & Martinez Arias, A. 2014. Wnt/β-catenin and FGF signalling direct the specification and maintenance of a neuromesodermal axial progenitor in ensembles of mouse embryonic stem cells. Development, 141, 4243–4253.

Tzouanacou, E., Wegener, A., Wymeersch, F., Wilson, V. & Nicolas, J. 2009. Redefining the progression of lineage segregations during mammalian embryogenesis by clonal analysis. Developmental Cell, 17, 365–76.

Verrier, L., Davidson, L., GierliŃski, M. & Storey, K. G. 2017. Generation, selection and transcriptomic profiling of human neuromesodermal and spinal cord progenitors in vitro. bioRxiv.

Watson, D. K., Reddy, E. P., Duesberg, P. H. & Papas, T. S. 1983. Nucleotide sequence analysis of the chicken c-myc gene reveals homologous and unique coding regions by comparison with the transforming gene of avian myelocytomatosis virus MC29, delta gag-myc. Proc Natl Acad Sci U S A, 80, 2146–50.

Weng, A. P., Millholland, J. M., Yashiro-Ohtani, Y., Arcangeli, M. L., Lau, A., Wai, C., Del Bianco, C., Rodriguez, C. G., Sai, H., Tobias, J., Li, Y., Wolfe, M. S., Shachaf, C., Felsher, D., Blacklow, S. C., Pear, W. S. & Aster, J. C. 2006. c-Myc is an important direct target of Notch1 in T-cell acute lymphoblastic leukemia/lymphoma. Genes & Development, 20, 2096–2109.

Wiedermann, G., Bone, R. A., Silva, J. C., Bjorklund, M., Murray, P. J. & Dale, J. K. 2015. A balance of positive and negative regulators determines the pace of the segmentation clock. Elife, 4, e05842.

Wilson, V., Olivera-Martinez, I. & Storey, K. G. 2009. Stem cells, signals and vertebrate body axis extension. Development, 136, 1591–604.

Wymeersch, F. J., Huang, Y., Blin, G., Cambray, N., Wilkie, R., Wong, F. C. K. & Wilson, V. 2016. Position-dependent plasticity of distinct progenitor types in the primitive streak. eLife, 5, e10042.

Yin, X., Giap, C., Lazo, J. S. & Prochownik, E. V. 2003. Low molecular weight inhibitors of Myc-Max interaction and function. Oncogene, 22, 6151–9.

Yu, P., Wilhelm, K., Dubrac, A., Tung, J. K., Alves, T. C., Fang, J. S., Xie, Y., Zhu, J., Chen, Z., De Smet, F., Zhang, J., Jin, S.-W., Sun, L., Sun, H., Kibbey, R. G., Hirschi, K. K., Hay, N., Carmeliet, P., Chittenden, T. W., Eichmann, A., Potente, M. & Simons, M. 2017. FGF-dependent metabolic control of vascular development. Nature, advance online publication.

Zeller, K. I., Jegga, A. G., Aronow, B. J., O’Donnell, K. A. & Dang, C. V. 2003. An integrated database of genes responsive to the Myc oncogenic transcription factor: identification of direct genomic targets. Genome Biol, 4, R69.

Zeller, K. I., Zhao, X., Lee, C. W., Chiu, K. P., Yao, F., Yustein, J. T., Ooi, H. S., Orlov, Y. L., Shahab, A., Yong, H. C., Fu, Y., Weng, Z., Kuznetsov, V. A., Sung, W. K., Ruan, Y., Dang, C. V. & Wei, C. L. 2006. Global mapping of c-Myc binding sites and target gene networks in human B cells. Proc Natl Acad Sci U S A, 103, 17834–9.

Zinin, N., Adameyko, I., Wilhelm, M., Fritz, N., Uhlen, P., Ernfors, P. & Henriksson, M. A. 2014. MYC proteins promote neuronal differentiation by controlling the mode of progenitor cell division. EMBO Rep, 15, 383–91.

